# Protein complex heterogeneity and topology revealed by electron capture charge reduction and surface induced dissociation

**DOI:** 10.1101/2024.03.07.583498

**Authors:** Jared B. Shaw, Sophie R. Harvey, Chen Du, Zhixin Xu, Regina M. Edgington, Eduardo Olmedillas, Erica Ollmann Saphire, Vicki H. Wysocki

## Abstract

We illustrate the utility of native mass spectrometry (nMS) combined with a fast, tunable gas-phase charge reduction, electron capture charge reduction (ECCR), for the characterization of protein complex topology and glycoprotein heterogeneity. ECCR efficiently reduces the charge states of tetradecameric GroEL, illustrating Orbitrap *m/z* measurements to greater than 100,000 *m/z*. For pentameric C-reactive protein and tetradecameric GroEL, our novel device combining ECCR with surface induced dissociation (SID) reduces the charge states and yields more topologically informative fragmentation. This is the first demonstration that ECCR yields more native-like SID fragmentation. ECCR also significantly improved mass and glycan heterogeneity measurements of heavily glycosylated SARS-CoV-2 spike protein trimer and thyroglobulin dimer. Protein glycosylation is important for structural and functional properties and plays essential roles in many biological processes. The immense heterogeneity in glycosylation sites and glycan structure poses significant analytical challenges that hinder a mechanistic understanding of the biological role of glycosylation. Without ECCR, average mass determination of glycoprotein complexes is available only through charge detection mass spectrometry or mass photometry. With narrow *m/z* selection windows followed by ECCR, multiple glycoform *m/z* values are apparent, providing quick global glycoform profiling and providing a future path for glycan localization on individual intact glycoforms.

## INTRODUCTION

Proteins and other biological macromolecules rarely work alone. The assembly and regulation of active cellular machinery are largely dependent on structure, which is modulated by subunit interactions, protein posttranslational modifications (PTMs), and ligand binding, among many other structure-modifying mechanisms. PTMs are covalent modifications of proteins and play a key role in numerous biological processes, affecting both structure and function. Full characterization of protein and nucleoprotein complexes with and without PTMs, especially large or complicated systems with many small and large partners, remains an analytical challenge. Of the potential modifications a protein can undergo, glycosylation is one of the most diverse and common PTMs (with up to one-fifth of proteins potentially being glycosylated)^1^, both with respect to the amino acids that can be modified and the modification structures. Monosaccharides can combine in a variety of ways with varying sequences, lengths, anomeric natures, and positions and branching points of linkages. In addition, the same site can be occupied by different glycosylation events on different copies of the same sequence. Characterization of glycosylation is essential as many therapeutic proteins are derived from endogenous glycoproteins, and a substantial number of currently approved protein therapeutics need to be properly glycosylated to exhibit optimal therapeutic efficacy.^2^

Native mass spectrometry (nMS) has emerged as a powerful tool for structural biology and enables the characterization of soluble and membrane protein complexes as well as heterogeneous assemblies, such as glycoprotein complexes, not readily suitable for complete high-resolution structural characterization.^3–6^ Although native MS does not provide atomic level structure information, it does have advantages in speed, sensitivity, selectivity, and the ability to simultaneously measure many components of a heterogeneous assembly or mixture.^7^ However, extreme heterogeneity in PTMs, assembly composition, and adduction of non-volatile salts can severely hinder mass determination and structure characterization by nMS. Several approaches have been presented in recent years to overcome these issues. These include charge manipulation approaches and charge detection mass spectrometry (CDMS), in which the charge and mass-to-charge ratio are simultaneously measured.^8–11^ CDMS overcomes the need for resolved charge states and provides insights into heterogeneous glycoproteins and large heterogeneous systems.^12–14^ Charge manipulation approaches typically involve reducing the charge either in solution or the gas-phase. This moves the charge state distribution to lower charges where the spacing between charge states is larger and enables resolution of heterogenous systems.^15,16^

Ion-neutral reactions and ion-ion reactions have been exploited to reduce the charge acquired by denatured proteins, native proteins, and protein complexes. Smith and co-workers demonstrated that gas-phase collisions between desolvated protein ions and ammonia, methylamines, and ethylamines resulted in effective charge reduction under denaturing and native conditions.^17–19^ Additionally, the pioneering work by McLuckey and Stephenson demonstrated the utility of ion-ion proton transfer reactions (PTR) for the study of ion chemistry as well as enhanced analytical capabilities for polypeptide characterization.^20–25^ In these experiments, a multiply charged polypeptide cation was reacted with a singly-charged reagent anion in an ion trapping device to generate a cation with reduced charge and a neutral reagent molecule.

PTR enabled analysis of mixtures of low, medium, and high molecular weight peptides and proteins by reducing spectral complexity introduced by multiple charging from electrospray ionization.^24^ Additionally, the development of “ion parking” techniques made it possible to inhibit ion-ion reaction rates via manipulation of ion velocities within ion trapping devices.^26^ This capability was demonstrated by concentration of ions originally present in a range of charge states into a selected charge state using PTR^26^ as well as inhibiting sequential electron transfer dissociation (ETD) reactions to enrich first-generation ETD product ions.^27^ Commercially, PTR and the related proton transfer charge reduction (PTCR) has been offered on various instrument platforms.^28–32^ Ogorzlek Loo et al. demonstrated charge reducing capabilities of corona discharge generated anions during the electrospray process.^33^ The Smith group utilized corona discharge or alpha-particle sources to generate anions used for charge manipulation via proton abstraction of electrosprayed proteins.^34,35^ Similarly, Bush and coworkers have used a glow-discharge source to generate anions for PTR with *m/z* -selected ions of native proteins and complexes to enable charge assignment and mass determination.^36^ Additionally, Bornschein and Ruotolo studied how charge reduction via corona discharge generated anions affected the collisional ejection of subunits from protein complexes.^37^ Sandoval and coworkers have recently demonstrated the utility of PTCR and gas-phase fractionation for the analysis of intact glycoproteins.^38^

Ion-ion and ion-electron reactions have predominantly been used to enable and enhance the characterization of peptides and denatured proteins by inducing covalent fragmentation for sequencing and PTM localization. However, several groups have utilized electron-based fragmentation to characterize higher-order protein structure.^39–44^ Electron-based fragmentation methods don’t generally impart sufficient energy to disrupt noncovalent interactions in addition to protein backbone fragmentation. Thus, ETD and its ion-electron reaction counterpart, electron capture dissociation (ECD), enable mapping of surface exposed and flexible regions of protein structure.^42–44^ Supplemental activation via collisions with, e.g., inert gas or infrared photons is needed to release additional fragments and obtain greater sequence coverage.^45–48^ Without supplemental activation to disrupt noncovalent interactions in larger proteins and protein complexes, ECD and ETD can yield abundant charge reduction without dissociation of the complementary N- and C-terminal peptide backbone cleavage product ions.

To date, non-dissociative electron capture or electron transfer events have generally been regarded as reaction by-products to be minimized to enhance the yield of peptide backbone cleavage products. However, some work on intact proteins and complexes has sought to utilize this by-product for the characterization of heterogeneous samples. Sobott and coworkers have demonstrated the utility of electron-based charge reduction to resolve heterogeneous oligomers of αβ-crystallin.^49^ Ujma and co-workers, and more recently Sobott and co-workers, have shown its utility to resolve AAV capsids,^50–52^ while Wysocki and coworkers have shown its ability to resolve fragments of AAV capsids. ^53^

Treated as an analytical tool, tunable electron transfer or electron capture charge reduction could be used to improve the apparent resolving power for large heterogeneous proteins and protein complexes by shifting ions to lower charge and higher m/z, as previously demonstrated using PTR. However, one challenge encountered with charge-reduced proteins and protein complexes is that they do not fragment as effectively as their high-charge counterparts. It is difficult to perform effective tandem mass spectrometry (MS/MS) via collisions with an inert gas, i.e., collision induced dissociation (CID)^54^ at energies accessible within practical limitations of the mass spectrometer hardware and electronics. Native protein complexes activated by CID dissociate via a monomer unfolding/complex restructuring mechanism. Typical products of CID are unfolded or elongated monomers with a disproportionate amount of charge and the corresponding (N-1) multimer; while these products confirm stoichiometry they provide limited information on overall complex topology/connectivity.^55^ Surface collisions, i.e., surface induced dissociation (SID), on the other hand, proceed via fast, high-energy deposition to yield more structurally informative subcomplexes, with the weakest protein complex interfaces cleaving at lower energues, producing products with compact structures and more proportionate charge partitioning.^56–62^ For example, a recent example of SID coupled with charge detection mass spectrometry shows extensive, structurally informative fragmentation of adeno-associated virus capsids (AAVs)^63^. To simplify SID integration into existing instruments, the Wysocki group has recently developed a simple yet novel split lens geometry SID device that was readily adapted and integrated into Q-IM-TOF, FTICR, and Orbitrap mass spectrometry platforms.^64^

Here, we combine electron capture charge reduction with SID, via a novel ExD cell^47,48^ designed for improved transmission of high *m/z* ions and facile integration of split lens SID electrodes at the exit of the ExD cell. The cell provides effective SID, extensive and tunable electron capture charge reduction (ECCR), or a combination of the two (ECCR-SID), with each of these modes illustrated here for large protein complexes. Tetradecameric GroEL (∼801 kDa) captured more than 50 electrons in a single pass through the cell and enabled observation of charge-reduced peaks at *m/z* greater than 100,000. Charge-reduced GroEL was also subjected to SID to determine whether the gas-phase charge-reduced precursors gave more native fragmentation. ECCR was also applied to heterogeneous glycoproteins, including a construct of the SARS-CoV-2 spike protein and thyroglobulin, better resolving the overlapping glycoform charge state distributions and yielding mass determination.

## RESULTS AND DISCUSSION

In previous work in our lab and others, when it has been desirable to obtain topological/connectivity information on protein complexes, low charge states have been used or the charge has been intentionally reduced. Charge reduction has typically been accomplished with solution additives (e.g., addition of the more basic triethylammonium acetate to the typical ammonium acetate electrolyte used for nanoelectrospray) or an alternative electrolyte (e.g., ethylene diammonium diacetate).^65–67^ While this accomplishes the needed charge reduction, solution additives, particularly triethylammonium acetate, often cause peak broadening, influencing mass accuracy. While adducts can be removed by collisional activation, this activation can restructure protein complexes, leading to non-native fragmentation of the complex and obscuring desirable topology information.^68,69^ There has thus been a strong need to accomplish charge reduction without adducting and peak broadening.

### Hybrid ExD-SID Cell

A prototype ExD cell was developed, in collaboration with e-MSion, with the goal of improving ease of use, transmission of a broad and high *m/z* range, and incorporation of electrodes to perform SID. Figure 1 shows schematics of the hybrid ExD-SID cell and integration of the cell into the Q Exactive UHMR Orbitrap mass spectrometer. Thick electrostatic lenses at each end of the standard ExD cell were replaced with quadrupole ion guides. The quadrupoles were constructed using 3/16-inch diameter precision ground stainless steel round rods with a rod radius to inscribed radius ratio of 1.14. DC voltages were supplied to the ExD portion of the cell using a commercial power supply and software from eMSion. The added SID electrode design and integration were similar to that previously described by Snyder and coworkers^64^. Briefly, the three-electrode split-lens design was integrated at the exit of the cell. Half of the split-lens is the SID surface electrode with a thickness of 3 mm. The other half is composed of the deflector and extractor electrodes with 1 mm thickness and separated by a 1 mm insulating spacer. The split lens aperture is 2 mm in diameter. The C-trap DC offset during injection steps was supplied by an external power supply to allow a larger range of voltages to be applied and therefore increasing the SID voltage range (defined as the difference in voltage between the bent flatapole and surface voltage), as previously described.^70^ C-Trap DC offset and SID device voltages were supplied via external power supplies (Ardara Technologies, Ardara, PA) and controlled by Tempus software (Ardara Technologies, Ardara, PA). The device works well as an SID device, as illustrated by fragmentation of the 50S ribosome subunit spectrum shown in Figure S1.

**Figure 1.**
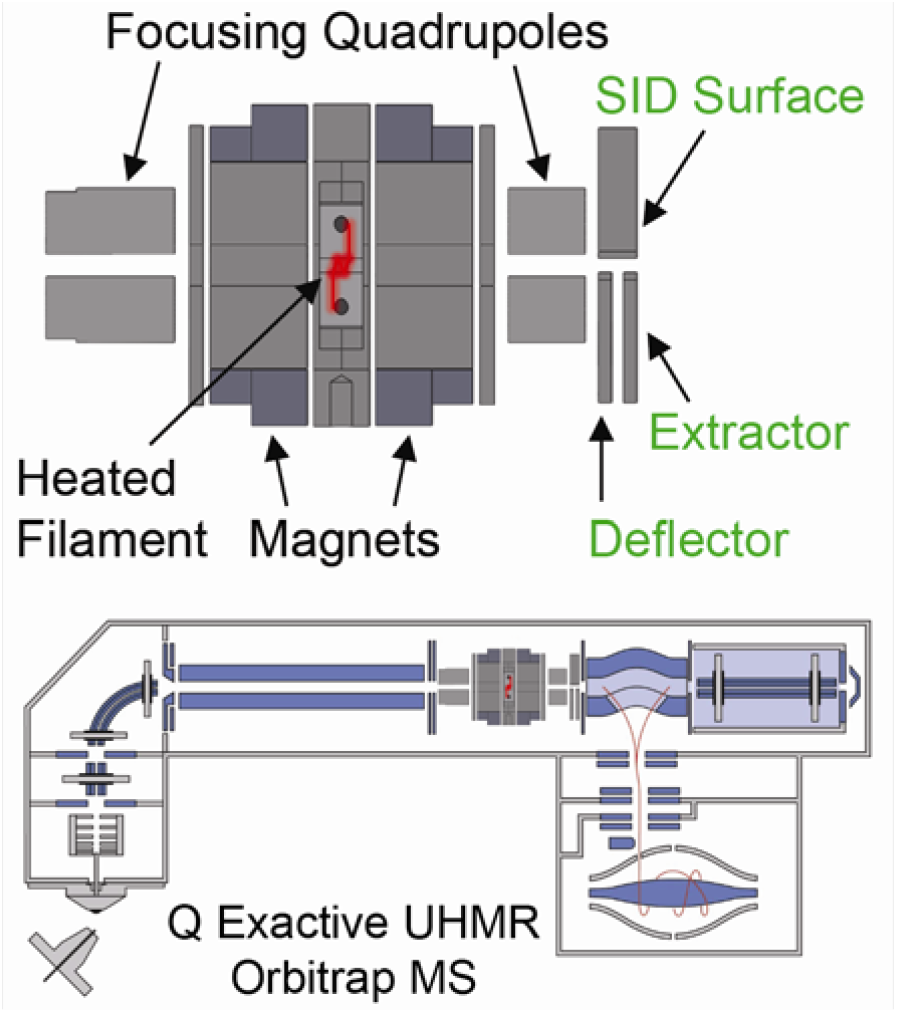
Schematics of the ExD-SID cell and integration of the cell into the Q Exactive UHMR Orbitrap mass spectrometer.

### Tunable Electron Capture Charge Reduction (ECCR)

The first goal of this work was to develop an approach for tunable gas phase charge reduction to improve the effective resolving power of heterogeneous protein complexes. By reducing the charge of the heterogeneous complex ions, the species detected are shifted to a greater *m/z* range thus increasing the Δ *m/z* for overlapping peaks, an approach that has been used by others to better resolve large or heterogeneous systems.^15,16,38,49,71–72^ For initial method development and proof of principle for ECCR, we first used GroEL,^51,74,75^ an ∼801 kDa homo-tetradecameric protein composed of two stacked seven-member rings that has been thoroughly studied by native mass spectrometry.^55,60,66,76,77^

GroEL prepared in 200 mM ammonium acetate and ionized by nano-electrospray ionization produced the “normal charge” mass spectrum shown at the top of Figure 2. The intensity-weighted average charge was 64+ with a charge state distribution, for peaks greater than 5% relative abundance, ranging from 67+ to 62+. Charge reduction was achieved via ion-electron reactions in which the multiply charged cations of GroEL captured multiple low-energy electrons while flying through the ExD-SID cell. For peptides and some proteins, ECD products (covalent bond cleavages) are the major expected products, but for large complexes, the major product is the intact non-covalent complex with multiple lower charge states. Detection of the intact mass of the charge-reduced species relies on preservation of GroEL subunit intramolecular and intermolecular non-covalent interactions. This is achieved using minimal collisional activation throughout the experiment.

**Figure 2.**
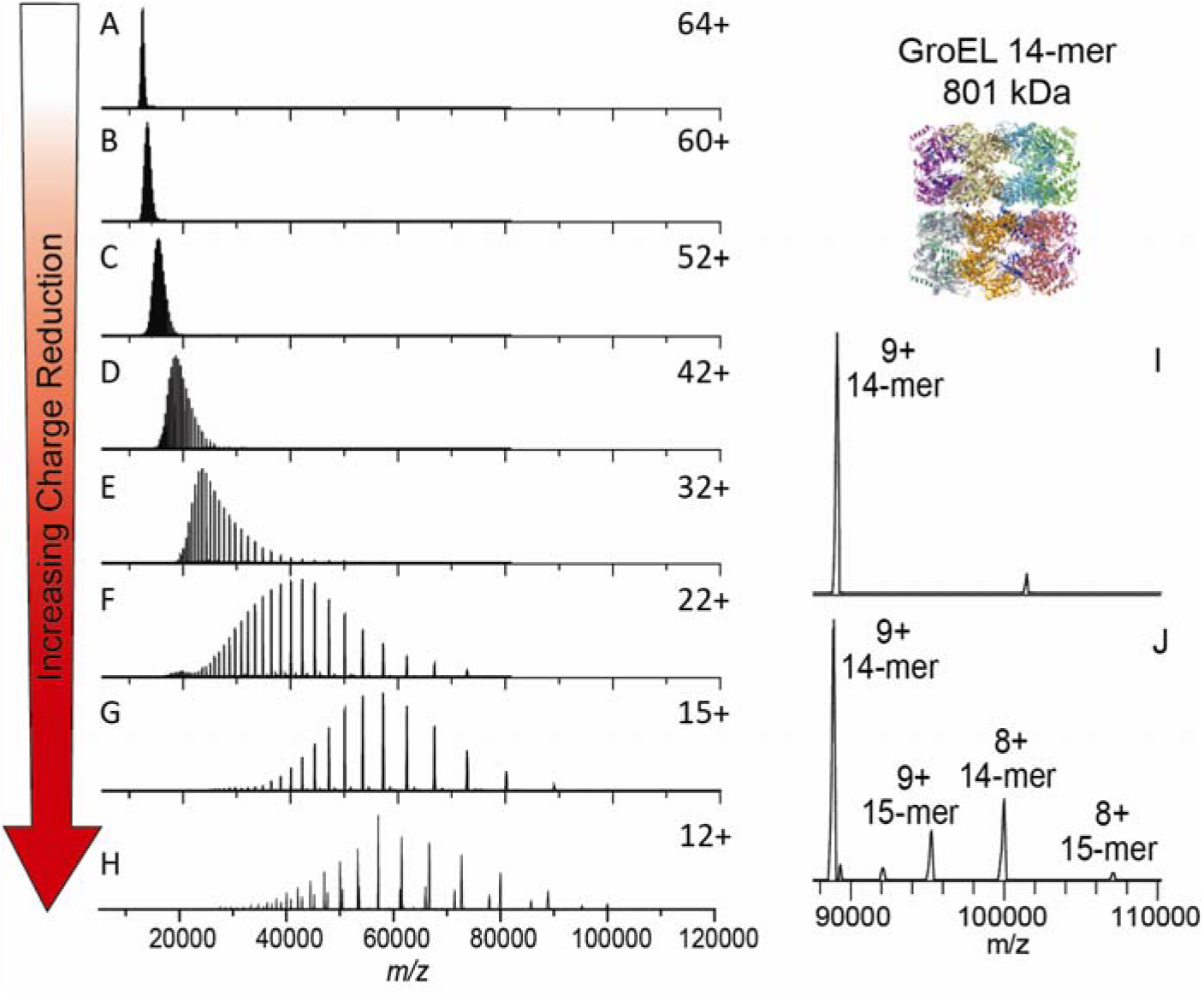
Electron capture charge reduction of GroEL. The top panel shows the mass spectrum of GroEL with the typical charge distribution generated by spraying from ammonium acetate (i.e., no gas phase charge reduction). Panels B-F show results from increases in electron capture charge reduction, with the device voltage that initiates ECCR increasing in two-volt steps (in-source trapping of –75V and trap gas setting of 7). Panel G is maximum ECCR obtained for this sample (see Table S1). Panel H is from a different sample from A-G and shows more extensive charge reduction with peaks up to 107K *m/z* ; more 15-mer of GroEL (additional, lower abundance peaks) was present in this sample, an older sample that had aged after refolding (in-source trapping of –100 V and trap gas setting of 8). Panel I is a zoom in on the high *m/z* range of panel H.

The extent of ECCR was modulated by adjusting the DC voltages applied to the center of the ExD device, namely the two magnetic lenses and filament holder (representative tuning conditions are given in Table S1 along with a labeled schematic of the ExD-SID cell). The voltages were incrementally increased to reduce the velocity of the ions in transit. The lower panels of Figure 2 show incremental increases in ECCR with maximum ECCR yielding a reduction in the average charge of GroEL from 64+ to 12+, with an average of 52 electrons captured at maximum ECCR. The width of the charge state distribution increased from 7 to 11 charge states with minimal ECCR and plateaued at approximately 22 for higher levels of ECCR before returning to 12 charge states (above 5% intensity) at maximum ECCR (Figure S2). This reflects the distribution of ion kinetic energies in the ion beam and the strong dependence of electron capture kinetics on the ion kinetic energy and charge state. All experiments shown here were performed using in-source trapping (−75 to -100 V) for desolvation.^78^ This process involves collisional activation followed by storage of the ions in the source region of the instrument for a few milliseconds. The momentum dampening and desolvation capabilities enabled by in-source trapping greatly increase transmission and sensitivity for high *m/z* ions, but some distribution in ion kinetic energy is unavoidable due to gas dynamics in the atmospheric pressure interface and spatial distribution of ions trapped in the injection flatapole. The extent of charge reduction exhibits a linear relationship when voltages are incrementally linearly increased (data shown in SI Figure S2). These results were reproduced and expanded upon in the recent ECCR results for GroEL shown by Sobott, Makarov, Fort, and co-workers. ^51,52^ Figure S3 demonstrates long-term stability and high reproducibility of ECCR for CRP, where it was observed the standard deviation in the average charge state after ECCR (± 1.9) is slightly higher than the standard deviation in the average charge state before ECCR (± 0.9), over 7 months of data acquisition.

### Coupling ECCR and Surface Induced Dissociation

Initial experiments with GroEL highlighted that ECCR is tunable, which could be advantageous for heterogeneous samples (as demonstrated with glycoproteins below) but we also wanted to determine whether ECCR affects the protein quaternary structure and if structurally relevant information could still be obtained from the protein complex after ECCR. This is especially important because many structural studies using nMS already employ charge-reducing conditions (typically with solution phase additives), as lower charge states can give more stable, native-like structures and fragmentation products.^65,79^

We used SID to probe the structure of protein complex ions after ECCR. It has been shown that protein complex dissociation via a surface collision can occur, depending on the complex, with reduced subunit or subcomplex unfolding compared to traditional inert gas collision induced dissociation, providing information consistent with the native structure.^59,80^ To investigate the effect of gas phase charge reduction on SID, we compared dissociation of the pentameric C-reactive protein complex (CRP) under normal charge (ammonium acetate), solution charge reduction, and gas phase charge reduction conditions (Figure 3). Analysis of CRP from 200 mM ammonium acetate yields a weighted average charge state of 23+ which will be referred to as normal charge for CRP (Figure 3A). SID of the entire charge state distribution of CRP at 40 V (energy range of 880 to 960 eV, determined by multiplying the SID voltage by the charge states observed above 5% relative intensity) produced primarily monomer and dimer, with lower levels of trimer and tetramer in agreement with previous studies (Figure 3B).^69,79^ We selected the full charge state distribution knowing that we won’t be able to select an individual charge state after ECCR, because the ExD-SID device is after the quadrupole mass filter.

**Figure 3.**
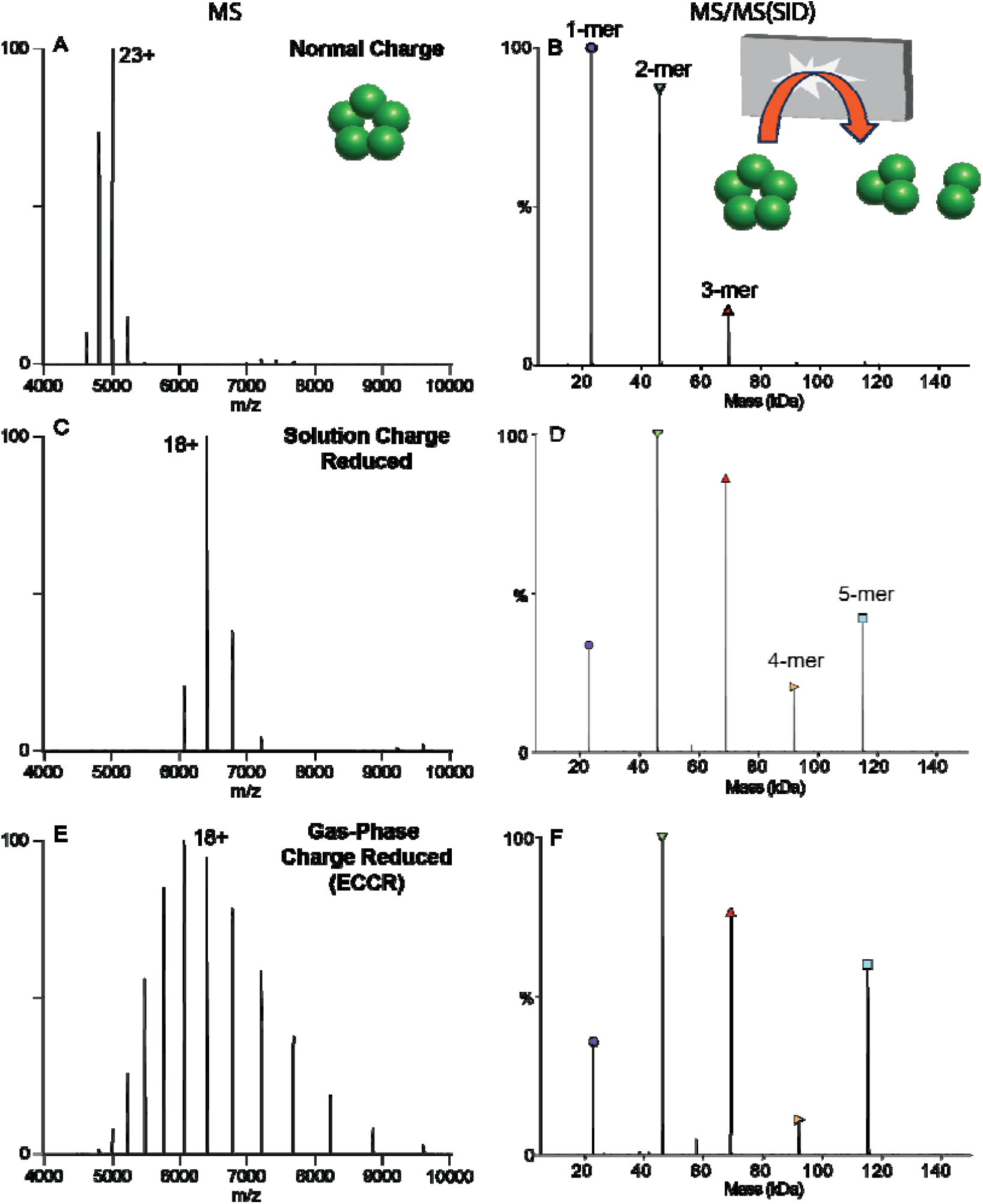
Comparison of C-reactive protein MS (left) and SID (right) patterns for A&B) normal charge (sprayed from ammonium acetate), C&D) solution charge reduced (sprayed from an 80:20 ratio of ammonium acetate to triethylammonium acetate) E&F) gas-phase charge reduced (ECCR). Acquired with in-source trapping -30 V, and SID voltage of 40 V (normal charge and gas-phase charge reduced) or 60 V (solution phase charge reduced).

For a cyclic complex like CRP, we expect all oligomeric states between monomer and tetramer at low SID energies, with relatively high abundance due to the equal interfaces between all subunits. Solution charge reduction of CRP (160 mM ammonium acetate and 40 mM triethylammonium acetate) yielded a charge state distribution with a weighted average charge of 18+ (Figure 3C). SID of this charge state distribution at 60 V (energy range of 1020-1140 eV) produced [monomer and tetramer] and [dimer and trimer] at high intensity (Figure 3D) consistent with the native cyclic structure and with previous SID studies of CRP.^59,69,81^ As noted above, charge reduced precursors are often chosen for native SID studies as the fragmentation has been observed to be more native-like, producing expected substructure units (e.g., dimers from dimer of dimers or trimer from dimer of trimers) and more of the higher-order oligomers due to decreased secondary fragmentation and/or unfolding.^69,79^

Finally, gas-phase charge reduction via ECCR of the normal charge distribution yielded a weighted average charge state of 18+ but with a broader charge state distribution (Figure 3E), which when subjected to SID at 40V (energy range of 560 to 880 eV) (Figure 3F) produced a similar SID spectrum to that obtained from solution-phase charge reduction (Figure 3D). Previous studies have shown solution charge reducing agents, such as TEAA, do not markedly change the gas-phase structure of precursor ions,^59^ but help stabilize noncovalently associated complexes and direct the course of gas-phase dissociation to give more native-like products.^67^ Therefore, the high degree of similarity between solution charge reduced and gas-phase charge reduced CRP indicates modulation of charge driven processes during dissociation rather than gas-phase structural changes resulting from charge neutralization via ECCR. These results show that ECCR-SID can probe the topology of protein complexes with dissociation occurring that is consistent with the solved structure.

We further investigated the ability of gas phase charge reduction (ECCR) to produce more native-like topology/connectivity information by comparing SID and ECCR followed by SID for the tetradecamer of GroEL. Figures 4A and B show the SID mass spectrum and charge deconvolved spectrum for normal charge (68+ weighted average charge) GroEL. The monomer of GroEL is by far the most abundant product ion with lower levels of dimer through 12mer. However, gas-phase charge reduction of normal charge GroEL (46+ weighted average charge) followed by SID (Figure 4C and D) yielded a significantly lower relative abundance of monomer and very abundant 7mer corresponding to dissociation at the interface of the two stacked 7mer rings. It should be noted that SID was performed at the same voltage in each case, corresponding to different SID energies because charge is reduced in the ExD-SID device and charge and electric field determine kinetic energy. To confirm that the kinetic energy difference did not cause the spectral differences, we lowered the kinetic energy for the normal charge GroEL and that did not yield a 7mer that is significantly increased as in Fig. 4 (see Figure S4).

**Figure 4.**
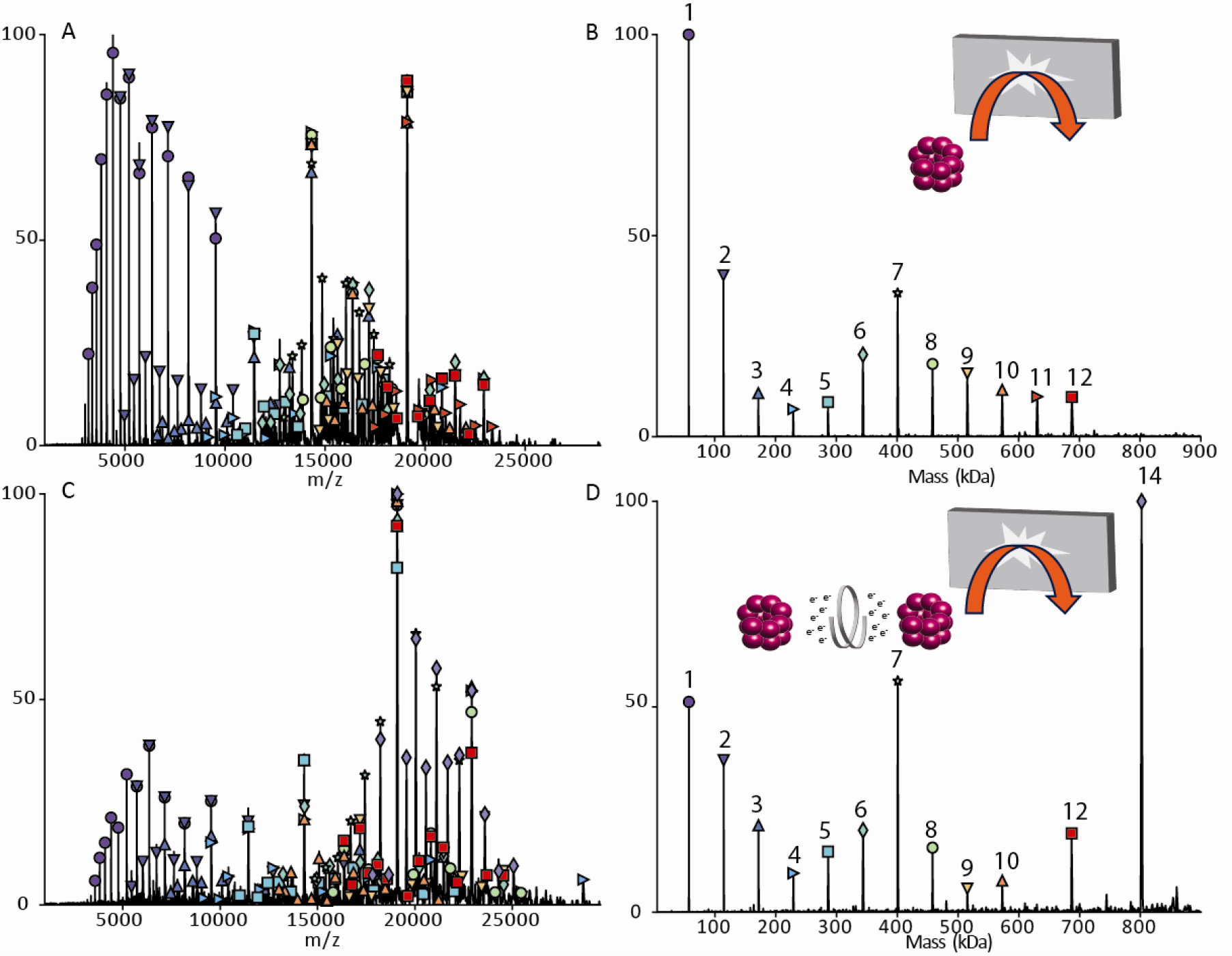
GroEL SID at 230 V.(A) Raw data for normal charge (68+ weighted average charge). (B) Deconvolved data for normal charge. (C) Raw data for gas-phase charge reduction (46+ weighted average charge). (D) Deconvolved data for gas-phase charge reduction. Acquired with in-source trapping set to –30 V.

### Application of ECCR to Glycoproteins

Posttranslational modifications pose significant challenges to mass spectrometric analysis at the intact protein and protein complex levels. For glycosylation, the challenge results from the immense complexity arising from glycosylation site occupancy (macroheterogeneity) and variations in the glycans at a given site (microheterogeneity). Electrospray ionization mass spectra of heterogeneously glycosylated proteins and protein complexes contain complex overlapping distributions of peaks resulting from the observation of multiple charge states and varying states of glycosylation.^38^ Heterogeneity is often sufficient to preclude mass determination due to the inability to resolve distinct isotopic and or charge state distributions. An example of this is shown in Figure 5 for VFLIP, a stable construct of the SARS-CoV-2 spike protein trimer with disulfide bonding between monomers and native-like glycosylation, which is of interest as an improved tool for diagnostics and vaccine design.^82^

**Figure 5.**
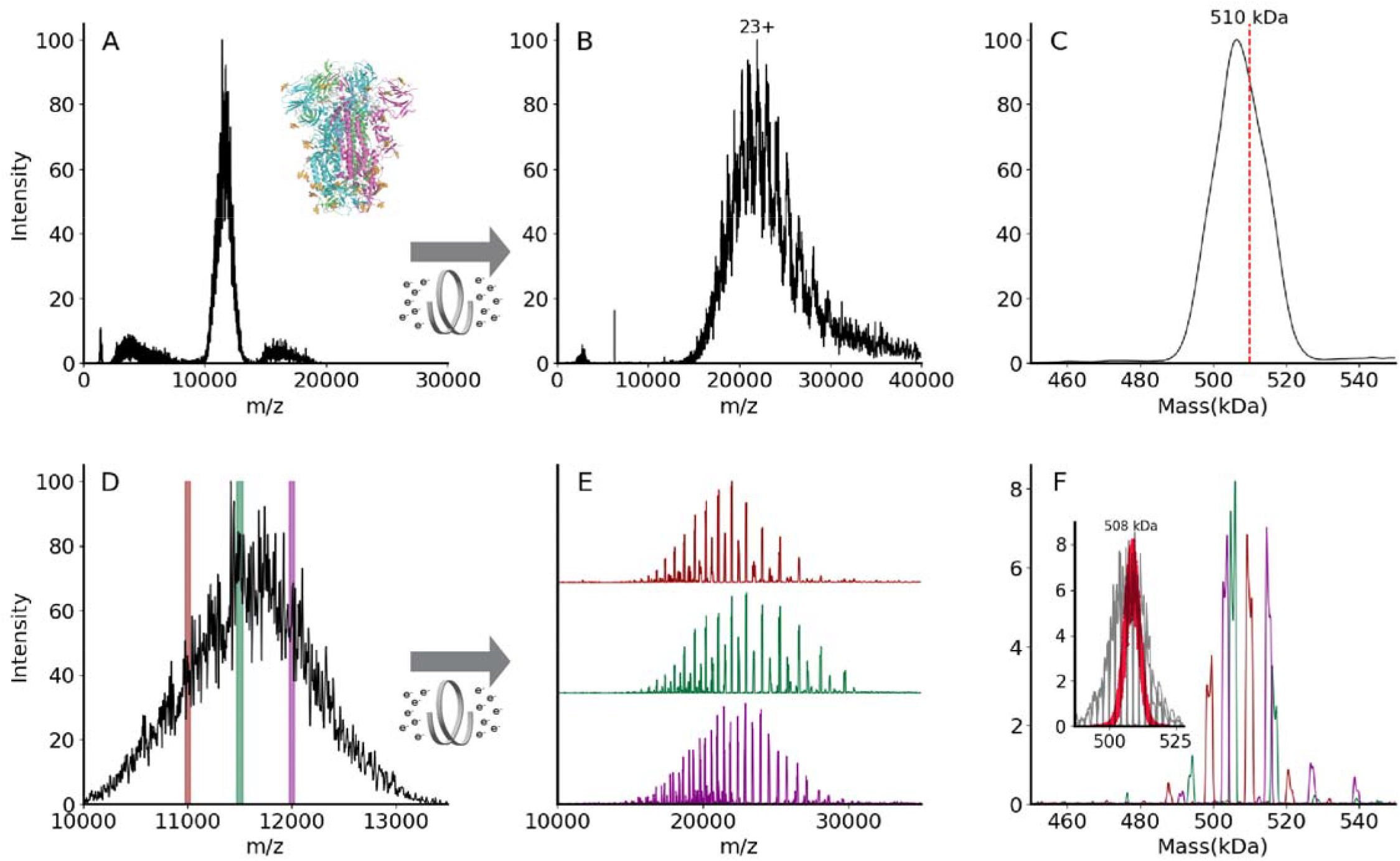
(A) An unresolved native mass spectrum of the heterogeneously glycosylated spike trimer VFLIP^82^. Inset shows the ribbon structure of SARS-CoV-2 spike trimer (PDB: 6×79). Three protomers are shown in pink, green, or blue, and N-glycans are shown in gold. (B) A charge state-resolved native mass spectrum of the spike trimer after electron capture charge reduction (ECCR, voltage 7V). (C) The average mass is 506 kDa deconvolved using UniDec parameters described in Methods^89^. The red dashed line indicates theoretical mass of 510 kDa. (D) Zoom in for spike trimer VFLIP from A with the positions of three narrow quadrupole selections shown. (E) Plots showing ECCR corresponding to the three narrow window isolations (10975-11025 (maroon); 11475-11525 (green); 11975-12025 (magenta), respectively. (F) Overlayed deconvolved mass spectra for data shown in panel E. The intensity was not normalized to better reflect the original intensity in panel D. The detected masses for first narrow window selection (maroon) are 499,645 Da, 509,355 Da, 520,689 Da; those for the second (green) are 494,331 Da, 504,835 Da, 506,070 Da, 516,276 Da, and 517,408 Da; those for the third selection (magenta) are 503,916 Da, 514,746, and 526,832 Da. Inset shows deconvolved masses from ECCR of 10 narrow isolation windows (50 *m/z* units wide, from 10,975-to 11,475 *m/z*) in gray with an overlay shown in red of the Monte Carlo simulations of theoretical masses based on the glycoproteomic data.

Figure 5A shows a broad and unresolved *m/z* distribution corresponding to the intact complex for which mass determination is not possible by traditional mass spectrometry methods. To resolve charge states of VFLIP, ECCR was applied to the entire *m/z* distribution of the intact complex, moderately reducing the charge in the gas phase. The resulting mass spectrum in Figure 5B shows resolved charge states of the VFLIP complex that enable mass determination. An average mass of 506 kDa was determined from UniDec, consistent with the expected mass based on sequence and known glycosylation profile (expected mass 510 kDa). The application of ECCR to the heterogeneous glycoprotein complex VFLIP illustrates the capability to determine mass and resolve sources of heterogeneity when traditional native MS approaches don’t provide that information. There are, however, alternative approaches that can be used for such heterogeneous samples. Mass photometry utilizes the linear relationship between light scattered from a single particle and its mass to measure masses of molecules.^83–85^ This instrument provides single molecule level studies in solution for proteins without labels.^86^ The mass was measured using MP and the mass distribution centered at 510 kDa (see Figure S5A). Charge detection mass spectrometry (CDMS)^8,9,87,11^ and the recently commercialized version of CDMS, Direct Mass Technology (DMT)^88^ for Orbitrap mass analyzers, enables simultaneous detection of *m/z* and charge (z) of analytes. This approach is particularly useful for mass determination of very large biomolecules, e.g., for heterogeneous spike proteins and megadalton size viral capsids, where heterogeneity in composition and incomplete desolvation yield very broad and unresolved spectra.^12,63,87^ CDMS was applied to a VFLIP spike protein and mass distributions centered at 513 kDa (HCD 0 V, no activation in the collision cell) and 509 kDa (HCD 200 V, activation in the collision cell to remove solvent and non-covalent adducts) were determined (see Figure S5B and C).

While we can determine the average mass with CDMS, ECCR, or mass photometry of the full charge state distribution, more detailed information on the different glycoforms present can be obtained using narrow window isolations and ECCR across the unresolved region, enabling better resolution of the different glycoforms (Figure 5). Three narrow window isolations across the unresolved charge state distribution (Figure 5D), followed by ECCR, give results illustrated in Figures 5E and F. ECCR of the narrow window isolation shows better-resolved peaks that can easily be deconvolved to gain more accurate mass measurements. Within each *m/z* selection window, 3-6 different mass species are observed (Figure 5F), reflecting the heterogeneity of the VFLIP glycosylation. For a protein complex with a lower total glycosylation mass, e.g., thyroglobulin, each *m/z* selection window results in a less complex distribution of glycoforms (see Figure S6) and narrow *m/z* selection windows reveal glycoforms. Overlaying the deconvolved mass peaks from multiple narrow window isolations for the spike protein (e.g., three shown in Fig. 5F reconstructs the original, unresolved VFLIP spike protein mass distribution, providing a glycan profile.

Three isolation windows, however, do not capture the sample’s full heterogeneity. To highlight the complexity and heterogeneity of this sample, we performed additional ECCR experiments on 10 adjacent narrow isolation windows (gray trace in the inset of Figure 5F) and compared the results to theoretical masses obtained by using a Monte Carlo simulation based on glycoproteomics data (red dots in the inset of Figure 5F).^90,91^ The proteomics-based Monte Carlo simulation implies a broad mass distribution for spike protein,^90,91^ with masses from 492 kDa to 522 kDa, with an average mass for spike protein of 508 kDa. However, the native MS ECCR results (gray) reflect a broader, more heterogeneous distribution. The broad distribution is like that reported by Jarrold, Clemmer, and Robinson by CDMS,^14^ but our ECCR results and proteomics-based Monte Carlo simulation results center at around the same mass rather than at different masses and distinct glycoform masses are available from the *m/z* window slicing and ECCR. There are several possible reasons that the width of the proteomics-based simulations and the native MS distribution might differ. 1) The difference may be caused, at least partially, by the different cell lines used for VFLIP samples (Chinese hamster ovary cell line) compared to those used for the glycoproteomics studies (Human embryonic kidney 293 cell line).^14,90,91^ 2) The glycoproteomics data (digested proteins) may not provide information on biological glycan crosstalk, where the glycan type at one site may influence the glycosylation on another site. 3) The proteomics results may not capture all possible glycopeptides because of stochastic data collection, chromatography issues, and/or dynamic range issues. Based on our results, we suggest that this native MS method involving narrow-window *m/z* selection coupled to ECCR could be utilized as a quick screen for glycan complexity under different expression conditions for therapeutic proteins or various variants of concern in infectious diseases. This approach could be coupled with in-depth glycoproteomic studies (top-down d/or bottom-up) when the variable identities at each glycosylation site are required.

## CONCLUSIONS

In this study, a prototype device combining ion-electron reactions and surface induced dissociation was developed and incorporated into a commercial ultra-high mass range Q-Orbitrap mass spectrometer for the characterization of native protein complexes. The new device enabled efficient and tunable gas-phase electron capture charge reduction (ECCR), as demonstrated for GroEL by enabling detection of 8+ tetradecamer and pentadecamer at greater than 100,000 *m/z*. Our current results (https://www.biorxiv.org/content/10.1101/2024.03.07.583498v1) ^74,75^ are in strong agreement with the related work of Sobott and coworkers.^51,52^ We use the acronym ECCR to describe this gas phase charge transfer^53,74,75^ and agree with Sobott and coworkers who adopted the phrase and suggest ^52^ that it should be used when investigators use electron capture to induce charge reduction (in contrast to using the phrase electron capture dissociation, which describes experiments where fragments resulting from covalent cleavage are the intended products). Furthermore, we demonstrated that gas-phase charge reduction produces charge states that give native-like surface induced dissociation fragments for C-reactive protein and GroEL. ECCR also yields mass spectra that enable mass determination and better resolve glycan heterogeneity in the stabilized VFLIP spike and thyroglobulin glycoproteins. ECCR is a rapid and tunable gas-phase charge reduction technique that is a fast alternative, or complementary, approach to charge detection mass spectrometry techniques for very large and heterogeneous biomolecular assemblies and is expected to enhance future MS-based structural biology approaches. The native MS glycoform distribution revealed by ECCR captures greater heterogeneity than suggested by Monte Carlo simulations based on proteomics data. Work is in progress that expands this approach to SARS-CoV-2 spike proteins from multiple variants of concern and other structural biology problems.

## SUPPORTING INFORMATION

The supporting information is available free of charge at

- Figures showing SID of 50S ribosome, ECCR-SID device schematic and typical voltages, characteristics of GroEL charge reduction, GroEL SID, mass photometry of VFLIP spike protein trimer, ECCR of thyroglobulin, reproducibility of ECCR for C-reactive protein, and experimental methods.

## Author Contributions

J.B.S. and S.R.H. contributed equally to the preparation of the manuscript.

## Notes

J.B.S. is a former employed by e-MSion, Inc. V.H.W. and S.R.H. hold intellectual property for SID.

## ACKNOWLEDGEMENTS

The authors thank Yury Vasil’ev and Joe Beckman (formerly of e-MSion, Inc.) for helpful conversations on the ExD device, Dalton Snyder (OSU) for helpful conversations on SID device geometry and tuning, Andrew Arslanian (OSU) for helping to collect the CDMS spectra in the supporting information, Jacelyn Greenwald (OSU) for refolding the GroEL samples, and Steffen Lindert and his group members for helpful suggestions on the Monte Carlo simulations. The authors gratefully acknowledge funding from the National Institutes of Health awarded to J.B.S (R43GM140749), E.O.S. (P01AI165072), and V.H.W (P41GM128577 and RM1GM149374). J.B.S gratefully acknowledges funding from the Nebraska Center for Integrated Biomolecular Communication (P20GM113126).

## TOC GRAPHIC

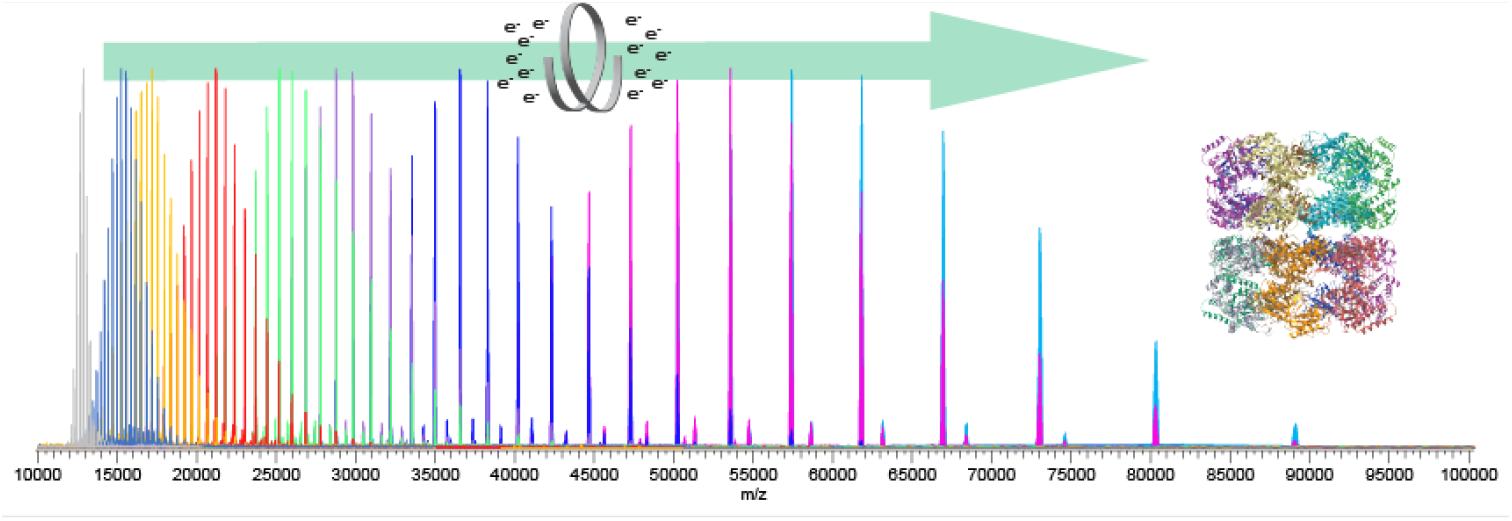

## SYNOPSIS

Electron capture charge reduction and surface induced dissociation enable more native-like fragmentation and define heterogeneity in glycosylated protein complexes.

## REFERENCES

(1) Khoury, G. A.; Baliban, R. C.; Floudas, C. A. Proteome-Wide Post-Translational Modification Statistics: Frequency Analysis and Curation of the Swiss-Prot Database. Sci Rep 2011, 1 (1), 90. 10.1038/srep00090.

(2) Solá, R. J.; Griebenow, K. Glycosylation of Therapeutic Proteins: An Effective Strategy to Optimize Efficacy. BioDrugs 2010, 24 (1), 9–21. 10.2165/11530550-000000000-00000.

(3) Heck, A. J. R. Native Mass Spectrometry: A Bridge between Interactomics and Structural Biology. Nat. Meth. 2008, 5 (11), 927–933. 10.1038/nmeth.1265.

(4) Barrera, N. P.; Robinson, C. V. Advances in the Mass Spectrometry of Membrane Proteins: From Individual Proteins to Intact Complexes. Annu. Rev. Biochem. 2011, 80 (1), 247–271. 10.1146/annurev-biochem-062309-093307.

(5) Allison, T. M.; Reading, E.; Liko, I.; Baldwin, A. J.; Laganowsky, A.; Robinson, C. V. Quantifying the Stabilizing Effects of Protein–Ligand Interactions in the Gas Phase. Nat Commun 2015, 6 (1), 8551. 10.1038/ncomms9551.

(6) Uetrecht, C.; Barbu, I. M.; Shoemaker, G. K.; van Duijn, E.; Heck, A. J. R. Interrogating Viral Capsid Assembly with Ion Mobility–Mass Spectrometry. Nature Chem 2011, 3 (2), 126–132. 10.1038/nchem.947.

(7) Lössl, P.; Snijder, J.; Heck, A. J. R. Boundaries of Mass Resolution in Native Mass Spectrometry. J. Am. Soc. Mass Spectrom. 2014. 10.1007/s13361-014-0874-3.

(8) Kafader, J. O.; Melani, R. D.; Durbin, K. R.; Ikwuagwu, B.; Early, B. P.; Fellers, R. T.; Beu, S. C.; Zabrouskov, V.; Makarov, A. A.; Maze, J. T.; Shinholt, D. L.; Yip, P. F.; Tullman-Ercek, D.; Senko, M. W.; Compton, P. D.; Kelleher, N. L. Multiplexed Mass Spectrometry of Individual Ions Improves Measurement of Proteoforms and Their Complexes. Nat Methods 2020, 17 (4), 391–394. 10.1038/s41592-020-0764-5.

(9) Wörner, T. P.; Snijder, J.; Bennett, A.; Agbandje-McKenna, M.; Makarov, A. A.; Heck, A. J. R. Resolving Heterogeneous Macromolecular Assemblies by Orbitrap-Based Single-Particle Charge Detection Mass Spectrometry. Nat Methods 2020, 17 (4), 395–398. 10.1038/s41592-020-0770-7.

(10) Harper, C. C.; Miller, Z. M.; Williams, E. R. Combined Multiharmonic Frequency Analysis for Improved Dynamic Energy Measurements and Accuracy in Charge Detection Mass Spectrometry. Anal. Chem. 2023, 95 (45), 16659–16667. 10.1021/acs.analchem.3c03160.

(11) Jarrold, M. F. Single-Ion Mass Spectrometry for Heterogeneous and High Molecular Weight Samples. J. Am. Chem. Soc. 2024, 146 (9), 5749–5758. 10.1021/jacs.3c08139.

(12) Stiving, A. Q.; Foreman, D. J.; VanAernum, Z. L.; Durr, E.; Wang, S.; Vlasak, J.; Galli, J.; Kafader, J. O.; Li, X.; Schuessler, H.; Richardson, D. Dissecting the Heterogeneous Glycan Profiles of Recombinant Coronavirus Spike Proteins with Individual Ion Mass Spectrometry. 2022. 10.26434/chemrxiv-2022-9vnch.

(13) Jarrold, M. F. Applications of Charge Detection Mass Spectrometry in Molecular Biology and Biotechnology. Chem. Rev. 2022, 122 (8), 7415–7441. 10.1021/acs.chemrev.1c00377.

(14) Miller, L. M.; Barnes, L. F.; Raab, S. A.; Draper, B. E.; El-Baba, T. J.; Lutomski, C. A.; Robinson, C. V.; Clemmer, D. E.; Jarrold, M. F. Heterogeneity of Glycan Processing on Trimeric SARS-CoV-2 Spike Protein Revealed by Charge Detection Mass Spectrometry. J Am Chem Soc 2021, 143 (10), 3959–3966. 10.1021/jacs.1c00353.

(15) Pacholarz, K. J.; Barran, P. E. Use of a Charge Reducing Agent to Enable Intact Mass Analysis of Cysteine-Linked Antibody-Drug-Conjugates by Native Mass Spectrometry. EuPA Open Proteomics 2016, 11, 23–27. 10.1016/j.euprot.2016.02.004.

(16) Bobst, C. E.; Sperry, J.; Friese, O. V.; Kaltashov, I. A. Simultaneous Evaluation of a Vaccine Component Microheterogeneity and Conformational Integrity Using Native Mass Spectrometry and Limited Charge Reduction. J. Am. Soc. Mass Spectrom. 2021, 32 (7), 1631–1637. 10.1021/jasms.1c00091.

(17) Loo, R. R. O.; Loo, J. A.; Udseth, H. R.; Fulton, J. L.; Smith, R. D. Protein Structural Effects in Gas Phase Ion/Molecule Reactions with Diethylamine. Rapid Commun. Mass Spectrom. 1992, 6 (3), 159–165. 10.1002/rcm.1290060302.

(18) Ogorzalek Loo, R. R.; Winger, B. E.; Smith, R. D. Proton Transfer Reaction Studies of Multiply Charged Proteins in a High Mass-to-Charge Ratio Quadrupole Mass Spectrometer. J. Am. Soc. Mass Spectrom. 1994, 5 (12), 1064–1071. 10.1016/1044-0305(94)85067-4.

(19) Ogorzalek Loo, R. R.; Smith, R. D. Investigation of the Gas-Phase Structure of Electrosprayed Proteins Using Ion-Molecule Reactions. J. Am. Soc. Mass Spectrom. 1994, 5 (4), 207–220. 10.1016/1044-0305(94)85011-9.

(20) McLuckey, S. A.; Stephenson, J. L.; Asano, K. G. Ion/Ion Proton-Transfer Kinetics:⍰ Implications for Analysis of Ions Derived from Electrospray of Protein Mixtures. Anal. Chem. 1998, 70 (6), 1198–1202. 10.1021/ac9710137.

(21) Stephenson, J. L.; McLuckey, S. A. Ion/Ion Reactions in the Gas Phase: Proton Transfer Reactions Involving Multiply-Charged Proteins. J. Am. Chem. Soc. 1996, 118 (31), 7390–7397. 10.1021/ja9611755.

(22) Stephenson, J. L.; McLuckey, S. A. Anion Effects on Storage and Resonance Ejection of High Mass-to-Charge Cations in Quadrupole Ion Trap Mass Spectrometry. Anal. Chem. 1997, 69 (18), 3760–3766. 10.1021/ac970399i.

(23) Stephenson, J. L.; McLuckey, S. A. Ion/Ion Proton Transfer Reactions for Protein Mixture Analysis. Anal. Chem. 1996, 68 (22), 4026–4032. 10.1021/ac9605657.

(24) Stephenson, J. L.; McLuckey, S. A. Ion/Ion Reactions for Oligopeptide Mixture Analysis: Application to Mixtures Comprised of 0.5–100 kDa Components. J. Am. Soc. Mass Spectrom. 1998, 9 (6), 585–596. 10.1016/S1044-0305(98)00025-7.

(25) Stephenson, J. L.; McLuckey, S. A. Simplification of Product Ion Spectra Derived from Multiply Charged Parent Ions via Ion/Ion Chemistry. Anal. Chem. 1998, 70 (17), 3533–3544. 10.1021/ac9802832.

(26) McLuckey, S. A.; Reid, G. E.; Wells, J. M. Ion Parking during Ion/Ion Reactions in Electrodynamic Ion Traps. Anal. Chem. 2002, 74 (2), 336–346. 10.1021/ac0109671.

(27) Chrisman, P. A.; Pitteri, S. J.; McLuckey, S. A. Parallel Ion Parking:⍰ Improving Conversion of Parents to First-Generation Products in Electron Transfer Dissociation. Anal. Chem. 2005, 77 (10), 3411–3414. 10.1021/ac0503613.

(28) Hartmer, R.; Kaplan, D. A.; Gebhardt, C. R.; Ledertheil, T.; Brekenfeld, A. Multiple Ion/Ion Reactions in the 3D Ion Trap: Selective Reagent Anion Production for ETD and PTR from a Single Compound. International Journal of Mass Spectrometry 2008, 276 (2), 82–90. 10.1016/j.ijms.2008.05.002.

(29) Huguet, R.; Mullen, C.; Srzentić, K.; Greer, J. B.; Fellers, R. T.; Zabrouskov, V.; Syka, J. E. P.; Kelleher, N. L.; Fornelli, L. Proton Transfer Charge Reduction Enables High-Throughput Top-Down Analysis of Large Proteoforms. Anal. Chem. 2019, 91 (24), 15732–15739. 10.1021/acs.analchem.9b03925.

(30) Kline, J. T.; Mullen, C.; Durbin, K. R.; Oates, R. N.; Huguet, R.; Syka, J. E. P.; Fornelli, L. Sequential Ion–Ion Reactions for Enhanced Gas-Phase Sequencing of Large Intact Proteins in a Tribrid Orbitrap Mass Spectrometer. J. Am. Soc. Mass Spectrom. 2021, 32 (9), 2334–2345. 10.1021/jasms.1c00062.

(31) Bailey, A. O.; Huguet, R.; Mullen, C.; Syka, J. E. P.; Russell, W. K. Ion–Ion Charge Reduction Addresses Multiple Challenges Common to Denaturing Intact Mass Analysis. Anal. Chem. 2022, 94 (9), 3930–3938. 10.1021/acs.analchem.1c04973.

(32) Sanders, J. D.; Mullen, C.; Watts, E.; Holden, D. D.; Syka, J. E. P.; Schwartz, J. C.; Brodbelt, J. S. Enhanced Sequence Coverage of Large Proteins by Combining Ultraviolet Photodissociation with Proton Transfer Reactions. Anal. Chem. 2020, 92 (1), 1041–1049. 10.1021/acs.analchem.9b04026.

(33) Ogorzalek Loo, R. R.; Udseth, H. R.; Smith, R. D. A New Approach for the Study of Gas-Phase Ion-Ion Reactions Using Electrospray Ionization. J Am Soc Mass Spectrom 1992, 3 (7), 695–705. 10.1016/1044-0305(92)87082-A.

(34) Scalf, M.; Westphall, M. S.; Krause, J.; Kaufman, S. L.; Smith, L. M. Controlling Charge States of Large Ions. Science 1999, 283 (5399), 194–197. 10.1126/science.283.5399.194.

(35) Ebeling, D. D.; Westphall, M. S.; Scalf, M.; Smith, L. M. Corona Discharge in Charge Reduction Electrospray Mass Spectrometry. Anal. Chem. 2000, 72 (21), 5158–5161. 10.1021/ac000559h.

(36) Laszlo, K. J.; Bush, M. F. Analysis of Native-Like Proteins and Protein Complexes Using Cation to Anion Proton Transfer Reactions (CAPTR). J. Am. Soc. Mass Spectrom. 2015, 26 (12), 2152–2161. 10.1007/s13361-015-1245-4.

(37) Bornschein, R. E.; Hyung, S.-J.; Ruotolo, B. T. Ion Mobility-Mass Spectrometry Reveals Conformational Changes in Charge Reduced Multiprotein Complexes. J. Am. Soc. Mass Spectrom. 2011, 22 (10). 10.1007/s13361-011-0204-y.

(38) Schachner, L. F.; Mullen, C.; Phung, W.; Hinkle, J. D.; Beardsley, M. I.; Bentley, T.; Day, P.; Tsai, C.; Sukumaran, S.; Baginski, T.; DiCara, D.; Agard, N.; Masureel, M.; Gober, J.; ElSohly, A.; Syka, J. E. P.; Huguet, R.; Marty, M. T.; Sandoval, W. Exposing the Molecular Heterogeneity of Glycosylated Biotherapeutics. bioRxiv May 11, 2023, p 2023.05.10.540271. 10.1101/2023.05.10.540271.

(39) Breuker, K.; Oh, H.; Horn, D. M.; Cerda, B. A.; McLafferty, F. W. Detailed Unfolding and Folding of Gaseous Ubiquitin Ions Characterized by Electron Capture Dissociation. J. Am. Chem. Soc. 2002, 124 (22), 6407–6420. 10.1021/ja012267j.

(40) Skinner, O. S.; McLafferty, F. W.; Breuker, K. How Ubiquitin Unfolds after Transfer into the Gas Phase. J. Am. Soc. Mass Spectrom. 2012, 23 (6), 1011–1014. 10.1007/s13361-012-0370-6.

(41) Harvey, S. Dissecting the Dynamic Conformations of the Metamorphic Protein Lymphotactin. The journal of physical chemistry. B 2014. 10.1021/jp504997k.

(42) Lermyte, F.; Sobott, F. Electron Transfer Dissociation Provides Higher-Order Structural Information of Native and Partially Unfolded Protein Complexes. PROTEOMICS 2015, 15 (16), 2813–2822. 10.1002/pmic.201400516.

(43) Li, H.; Nguyen, H. H.; Ogorzalek Loo, R. R.; Campuzano, I. D. G.; Loo, J. A. An Integrated Native Mass Spectrometry and Top-down Proteomics Method That Connects Sequence to Structure and Function of Macromolecular Complexes. Nat. Chem. 2018, 10 (2), 139–148. 10.1038/nchem.2908.

(44) Zhou, M.; Liu, W.; Shaw, J. B. Charge Movement and Structural Changes in the Gas-Phase Unfolding of Multimeric Protein Complexes Captured by Native Top-Down Mass Spectrometry. Anal. Chem. 2020, 92 (2), 1788–1795. 10.1021/acs.analchem.9b03469.

(45) Horn, D. M.; Ge, Y.; McLafferty, F. W. Activated Ion Electron Capture Dissociation for Mass Spectral Sequencing of Larger (42 kDa) Proteins. Anal. Chem. 2000, 72 (20), 4778–4784. 10.1021/ac000494i.

(46) Riley, N. M.; Westphall, M. S.; Coon, J. J. Sequencing Larger Intact Proteins (30-70 kDa) with Activated Ion Electron Transfer Dissociation. J. Am. Soc. Mass Spectrom. 2018, 29, 140–149. 10.1007/s13361-017-1808-7.

(47) Shaw, J. B.; Malhan, N.; Vasil’ev, Y. V.; Lopez, N. I.; Makarov, A.; Beckman, J. S.; Voinov, V. G. Sequencing Grade Tandem Mass Spectrometry for Top–Down Proteomics Using Hybrid Electron Capture Dissociation Methods in a Benchtop Orbitrap Mass Spectrometer. Anal. Chem. 2018, 90 (18), 10819–10827. 10.1021/acs.analchem.8b01901.

(48) Shaw, J. B.; Liu, W.; Vasil⍰ev, Y. V.; Bracken, C. C.; Malhan, N.; Guthals, A.; Beckman, J. S.; Voinov, V. G. Direct Determination of Antibody Chain Pairing by Top-down and Middle-down Mass Spectrometry Using Electron Capture Dissociation and Ultraviolet Photodissociation. Anal. Chem. 2020, 92 (1), 766–773. 10.1021/acs.analchem.9b03129.

(49) Lermyte, F.; Williams, J. P.; Brown, J. M.; Martin, E. M.; Sobott, F. Extensive Charge Reduction and Dissociation of Intact Protein Complexes Following Electron Transfer on a Quadrupole-Ion Mobility-Time-of-Flight MS. J. Am. Soc. Mass Spectrom. 2015, 26 (7), 1068–1076. 10.1007/s13361-015-1124-z.

(50) Ujma, J.; Giles, K.; Anderson, M.; Richardson, K. ENHANCED DECLUSTERING AND CHARGE-STRIPPING ENABLES MASS DETERMINATION OF AAVS IN TOF MS. 310509 MP 481, Proceedings of the 70th ASMS Conference on Mass Spectrometry and Allied Topics, Minneapolis, Minnesota. June5-9, 2022.

(51) Le Huray, K. I. P.; Woerner, T. P.; Reinhardt-Szyba, M.; Fort, K. L.; Sobott, F.; Makarov, A. A. To 200k m/z and beyond: Native Electron-Capture Charge Reduction Resolves Heterogeneous Signals in Large Biopharmaceutical Analytes, a New Orbitrap Record. 313520 (WOA pm 03:10), Proceedings of the 71st ASMS Conference on Mass Spectrometry and Allied Topics, Houston, Texas. June 4-8, 2023.

(52) Kyle I. P. Le Huray; Tobias P. Wörner; Tiago Moreira; Maria Reinhardt-Szyba; Paul W. A. Devine; Nicholas J. Bond; Kyle L. Fort; Alexander A. Makarov; Frank Sobott. To 200,000 m/z and beyond: Native Electron Capture Charge Reduction Mass Spectrometry Deconvolves Heterogeneous Signals in Large Biopharmaceutical Analytes. bioRxiv 2024, 2024.02.19.581059. 10.1101/2024.02.19.581059.

(53) Kostelic, M. M.; Du, C.; Wysocki, V. H. Architecture Of Adeno-Associated Viral Capsids With Surface-Induced Dissociation And Charge Detection Mass Spectrometry. 315406 (WP 233), Proceedings of the 71st ASMS Conference on Mass Spectrometry and Allied Topics, Houston, Texas. June 4-8, 2023.

(54) Lorenzen, K.; Versluis, C.; van Duijn, E.; van den Heuvel, R. H. H.; Heck, A. J. R. Optimizing Macromolecular Tandem Mass Spectrometry of Large Non-Covalent Complexes Using Heavy Collision Gases. International Journal of Mass Spectrometry 2007, 268 (2), 198–206. 10.1016/j.ijms.2007.06.012.

(55) Sobott, F.; Robinson, C. V. Characterising Electrosprayed Biomolecules Using Tandem-MS—the Noncovalent GroEL Chaperonin Assembly. International Journal of Mass Spectrometry 2004, 236 (1), 25–32. 10.1016/j.ijms.2004.05.010.

(56) Jones, C. M.; Beardsley, R. L.; Galhena, A. S.; Dagan, S.; Cheng, G.; Wysocki, V. H. Symmetrical Gas-Phase Dissociation of Noncovalent Protein Complexes via Surface Collisions. J. Am. Chem. Soc. 2006, 128 (47), 15044–15045. 10.1021/ja064586m.

(57) Blackwell, A. E.; Dodds, E. D.; Bandarian, V.; Wysocki, V. H. Revealing the Quaternary Structure of a Heterogeneous Noncovalent Protein Complex through Surface-Induced Dissociation. Anal. Chem. 2011, 83 (8), 2862–2865. 10.1021/ac200452b.

(58) Zhou, M.; Huang, C.; Wysocki, V. H. Surface-Induced Dissociation of Ion Mobility-Separated Noncovalent Complexes in a Quadrupole/Time-of-Flight Mass Spectrometer. Anal. Chem. 2012, 84 (14), 6016–6023. 10.1021/ac300810u.

(59) Zhou, M.; Dagan, S.; Wysocki, V. H. Protein Subunits Released by Surface Collisions of Noncovalent Complexes: Nativelike Compact Structures Revealed by Ion Mobility Mass Spectrometry. Angew. Chem. Int. Ed. 2012, 51 (18), 4336–4339. 10.1002/anie.201108700.

(60) Zhou, M.; Jones, C. M.; Wysocki, V. H. Dissecting the Large Noncovalent Protein Complex GroEL with Surface-Induced Dissociation and Ion Mobility–Mass Spectrometry. Anal. Chem. 2013, 85 (17), 8262–8267. 10.1021/ac401497c.

(61) Harvey, S. Relative Interfacial Cleavage Energetics of Protein Complexes Revealed by Surface Collisions. Proceedings of the National Academy of Sciences of the United States of America 2019. 10.1073/pnas.1817632116.

(62) Snyder, D. T.; Harvey, S. R.; Wysocki, V. H. Surface-Induced Dissociation Mass Spectrometry as a Structural Biology Tool. Chemical Reviews 2021. 10.1021/acs.chemrev.1c00309.

(63) Du, C.; Cleary, S. P.; Kostelic, M. M.; Jones, B. J.; Kafader, J. O.; Wysocki, V. H. Combining Surface-Induced Dissociation and Charge Detection Mass Spectrometry to Reveal the Native Topology of Heterogeneous Protein Complexes. Anal. Chem. 2023, 95 (37), 13889–13896. 10.1021/acs.analchem.3c02185.

(64) Snyder, D. T.; Panczyk, E. M.; Somogyi, A.; Kaplan, D. A.; Wysocki, V. Simple and Minimally Invasive SID Devices for Native Mass Spectrometry. Anal. Chem. 2020, 92 (16), 11195–11203. 10.1021/acs.analchem.0c01657.

(65) Hall, Z.; Politis, A.; Bush, M. F.; Smith, L. J.; Robinson, C. V. Charge-State Dependent Compaction and Dissociation of Protein Complexes: Insights from Ion Mobility and Molecular Dynamics. J. Am. Chem. Soc. 2012, 134 (7), 3429–3438. 10.1021/ja2096859.

(66) Dyachenko, A.; Gruber, R.; Shimon, L.; Horovitz, A.; Sharon, M. Allosteric Mechanisms Can Be Distinguished Using Structural Mass Spectrometry. PNAS 2013, 110 (18), 7235–7239. 10.1073/pnas.1302395110.

(67) Pagel, K.; Hyung, S.-J.; Ruotolo, B. T.; Robinson, C. V. Alternate Dissociation Pathways Identified in Charge-Reduced Protein Complex Ions. Anal. Chem. 2010, 82 (12), 5363–5372. 10.1021/ac101121r.

(68) Quintyn, R. S.; Zhou, M.; Yan, J.; Wysocki, V. H. Surface-Induced Dissociation Mass Spectra as a Tool for Distinguishing Different Structural Forms of Gas-Phase Multimeric Protein Complexes. Anal. Chem. 2015, 87 (23), 11879–11886. 10.1021/acs.analchem.5b03441.

(69) Harvey, S.; VanAernum, Z.; Wysocki, V. Surface-Induced Dissociation of Anionic vs Cationic Native-like Protein Complexes. 2021. 10.26434/chemrxiv.13547837.v1.

(70) VanAernum, Z. L.; Gilbert, J. D.; Belov, M. E.; Makarov, A. A.; Horning, S. R.; Wysocki, V. H. Surface-Induced Dissociation of Noncovalent Protein Complexes in an Extended Mass Range Orbitrap Mass Spectrometer. Anal. Chem. 2019, 91 (5), 3611–3618. 10.1021/acs.analchem.8b05605.

(71) Yang, Y.; Ivanov, D. G.; Kaltashov, I. A. The Challenge of Structural Heterogeneity in the Native Mass Spectrometry Studies of the SARS-CoV-2 Spike Protein Interactions with Its Host Cell-Surface Receptor. Anal Bioanal Chem 2021, 413 (29), 7205–7214. 10.1007/s00216-021-03601-3.

(72) Yang, Y.; Niu, C.; Bobst, C. E.; Kaltashov, I. A. Charge Manipulation Using Solution and Gas-Phase Chemistry to Facilitate Analysis of Highly Heterogeneous Protein Complexes in Native Mass Spectrometry. Anal. Chem. 2021, 93 (7), 3337–3342. 10.1021/acs.analchem.0c05249.

(73) Chen, Q.; Dai, R.; Yao, X.; Chaihu, L.; Tong, W.; Huang, Y.; Wang, G. Improving Accuracy in Mass Spectrometry-Based Mass Determination of Intact Heterogeneous Protein Utilizing the Universal Benefits of Charge Reduction and Alternative Gas-Phase Reactions. Anal. Chem. 2022, 94 (40), 13869–13878. 10.1021/acs.analchem.2c02586.

(74) Vasil’ev, Y. V.; Du, C.; Harvey, S. R.; Snyder, D. T.; Wysocki, V. H.; Shaw, J. B. Native Top-Down Characterization of Protein Complexes with a Hybrid ECD-SID Device. 311519 (WP176), Proceedings of the 70th ASMS Conference on Mass Spectrometry and Allied Topics, Minneapolis, Minnesota. June5-9, 2022.

(75) Shaw, J. B.; Harvey, S. R.; Du, C.; Wysocki, V. H. Protein Complex Heterogeneity and Structure Revealed by Native MS with ECCR and SID. 316163 (WOA pm 03:50), Proceedings of the 71st ASMS Conference on Mass Spectrometry and Allied Topics, Houston, Texas. June 4-8, 2023.

(76) Walker, T.; Sun, H. M.; Gunnels, T.; Wysocki, V.; Laganowsky, A.; Rye, H.; Russell, D. Dissecting the Thermodynamics of ATP Binding to GroEL One Nucleotide at a Time. ACS Cent. Sci. 2023, 9 (3), 466–475. 10.1021/acscentsci.2c01065.

(77) Walker, T. E.; Shirzadeh, M.; Sun, H. M.; McCabe, J. W.; Roth, A.; Moghadamchargari, Z.; Clemmer, D. E.; Laganowsky, A.; Rye, H.; Russell, D. H. Temperature Regulates Stability, Ligand Binding (Mg2+ and ATP), and Stoichiometry of GroEL-GroES Complexes. J Am Chem Soc 2022, 144 (6), 2667–2678. 10.1021/jacs.1c11341.

(78) Fort, K. L.; van de Waterbeemd, M.; Boll, D.; Reinhardt-Szyba, M.; Belov, M. E.; Sasaki, E.; Zschoche, R.; Hilvert, D.; Makarov, A. A.; Heck, A. J. R. Expanding the Structural Analysis Capabilities on an Orbitrap-Based Mass Spectrometer for Large Macromolecular Complexes. Analyst 2018, 143 (1), 100–105. 10.1039/C7AN01629H.

(79) Zhou, M.; Dagan, S.; Wysocki, V. H. Impact of Charge State on Gas-Phase Behaviors of Noncovalent Protein Complexes in Collision Induced Dissociation and Surface Induced Dissociation. Analyst 2013, 138 (5), 1353–1362. 10.1039/C2AN36525A.

(80) Harvey, S. R.; Seffernick, J. T.; Quintyn, R. S.; Song, Y.; Ju, Y.; Yan, J.; Sahasrabuddhe, A. N.; Norris, A.; Zhou, M.; Behrman, E. J. Relative Interfacial Cleavage Energetics of Protein Complexes Revealed by Surface Collisions. Proceedings of the National Academy of Sciences 2019, 116 (17), 8143–8148.

(81) Busch, F.; VanAernum, Z. L.; Ju, Y.; Yan, J.; Gilbert, J. D.; Quintyn, R. S.; Bern, M.; Wysocki, V. H. Localization of Protein Complex Bound Ligands by Surface-Induced Dissociation High-Resolution Mass Spectrometry. Anal. Chem. 2018, 90 (21), 12796–12801. 10.1021/acs.analchem.8b03263.

(82) Olmedillas, E.; Mann, C. J.; Peng, W.; Wang, Y.-T.; Avalos, R. D.; Bedinger, D.; Valentine, K.; Shafee, N.; Schendel, S. L.; Yuan, M.; Lang, G.; Rouet, R.; Christ, D.; Jiang, W.; Wilson, I. A.; Germann, T.; Shresta, S.; Snijder, J.; Saphire, E. O. Structure-Based Design of a Highly Stable, Covalently-Linked SARS-CoV-2 Spike Trimer with Improved Structural Properties and Immunogenicity; preprint; Microbiology, 2021. 10.1101/2021.05.06.441046.

(83) Cole, D.; Young, G.; Weigel, A.; Sebesta, A.; Kukura, P. Label-Free Single-Molecule Imaging with Numerical-Aperture-Shaped Interferometric Scattering Microscopy. ACS Photonics 2017, 4 (2), 211–216. 10.1021/acsphotonics.6b00912.

(84) Young, G.; Hundt, N.; Cole, D.; Fineberg, A.; Andrecka, J.; Tyler, A.; Olerinyova, A.; Ansari, A.; Marklund, E. G.; Collier, M. P.; Chandler, S. A.; Tkachenko, O.; Allen, J.; Crispin, M.; Billington, N.; Takagi, Y.; Sellers, J. R.; Eichmann, C.; Selenko, P.; Frey, L.; Riek, R.; Galpin, M. R.; Struwe, W. B.; Benesch, J. L. P.; Kukura, P. Quantitative Mass Imaging of Single Molecules. Science 2018, 360 (6387), 423–427. 10.1126/science.aar5839.

(85) Wu, D.; Piszczek, G. Standard Protocol for Mass Photometry Experiments. Eur Biophys J 2021, 50 (3–4), 403–409. 10.1007/s00249-021-01513-9.

(86) Sonn-Segev, A.; Belacic, K.; Bodrug, T.; Young, G.; VanderLinden, R. T.; Schulman, B. A.; Schimpf, J.; Friedrich, T.; Dip, P. V.; Schwartz, T. U.; Bauer, B.; Peters, J.-M.; Struwe, W. B.; Benesch, J. L. P.; Brown, N. G.; Haselbach, D.; Kukura, P. Quantifying the Heterogeneity of Macromolecular Machines by Mass Photometry. Nat Commun 2020, 11 (1), 1772. 10.1038/s41467-020-15642-w.

(87) Keifer, D. Z.; Pierson, E. E.; Jarrold, M. F. Charge Detection Mass Spectrometry: Weighing Heavier Things. Analyst 2017, 142 (10), 1654–1671. 10.1039/C7AN00277G.

(88) Kafader, J. O.; Beu, S. C.; Early, B. P.; Melani, R. D.; Durbin, K. R.; Zabrouskov, V.; Makarov, A. A.; Maze, J. T.; Shinholt, D. L.; Yip, P. F.; Kelleher, N. L.; Compton, P. D.; Senko, M. W. STORI Plots Enable Accurate Tracking of Individual Ion Signals. J Am Soc Mass Spectrom 2019, 30 (11), 2200–2203. 10.1007/s13361-019-02309-0.

(89) Marty, M. T.; Baldwin, A. J.; Marklund, E. G.; Hochberg, G. K. A.; Benesch, J. L. P.; Robinson, C. V. Bayesian Deconvolution of Mass and Ion Mobility Spectra: From Binary Interactions to Polydisperse Ensembles. Anal. Chem. 2015, 87 (8), 4370–4376. 10.1021/acs.analchem.5b00140.

(90) Wang, D.; Zhou, B.; Keppel, T. R.; Solano, M.; Baudys, J.; Goldstein, J.; Finn, M. G.; Fan, X.; Chapman, A. P.; Bundy, J. L.; Woolfitt, A. R.; Osman, S. H.; Pirkle, J. L.; Wentworth, D. E.; Barr, J. R. N-Glycosylation Profiles of the SARS-CoV-2 Spike D614G Mutant and Its Ancestral Protein Characterized by Advanced Mass Spectrometry. Sci Rep 2021, 11 (1), 23561. 10.1038/s41598-021-02904-w.

(91) Sanda, M.; Morrison, L.; Goldman, R. N- and O-Glycosylation of the SARS-CoV-2 Spike Protein. Anal. Chem. 2021, 93 (4), 2003–2009. 10.1021/acs.analchem.0c03173.

