## Supplementary information for "Protein complex heterogeneity and topology revealed by electron capture charge reduction and surface induced dissociation"

### Supporting information:

### Protein complex heterogeneity and structure revealed by native mass spectrometry with electron capture charge reduction and surface induced dissociation

Jared Shaw,[1,†,*] Sophie R. Harvey,[2,†] Chen Du,[2,3] Zhixin Xu,[2,3] Regina M. Edgington,[3] Eduardo Olmedillas,[4] Erica Ollmann Saphire,[4,5] Vicki H. Wysocki[2,3,*]

[1]Department of Chemistry, University of Nebraska-Lincoln, Lincoln, NE 68588; [2]Native Mass Spectrometry Guided Structural Biology Center, The Ohio State University, Columbus, OH 43210; [3]Department of Chemistry and Biochemistry, The Ohio State University, Columbus, OH 43210; [4]Center for Vaccine Innovation, La Jolla Institute for Immunology, La Jolla, CA 92037; [5]Department of Medicine, University of California San Diego, La Jolla, CA 92037

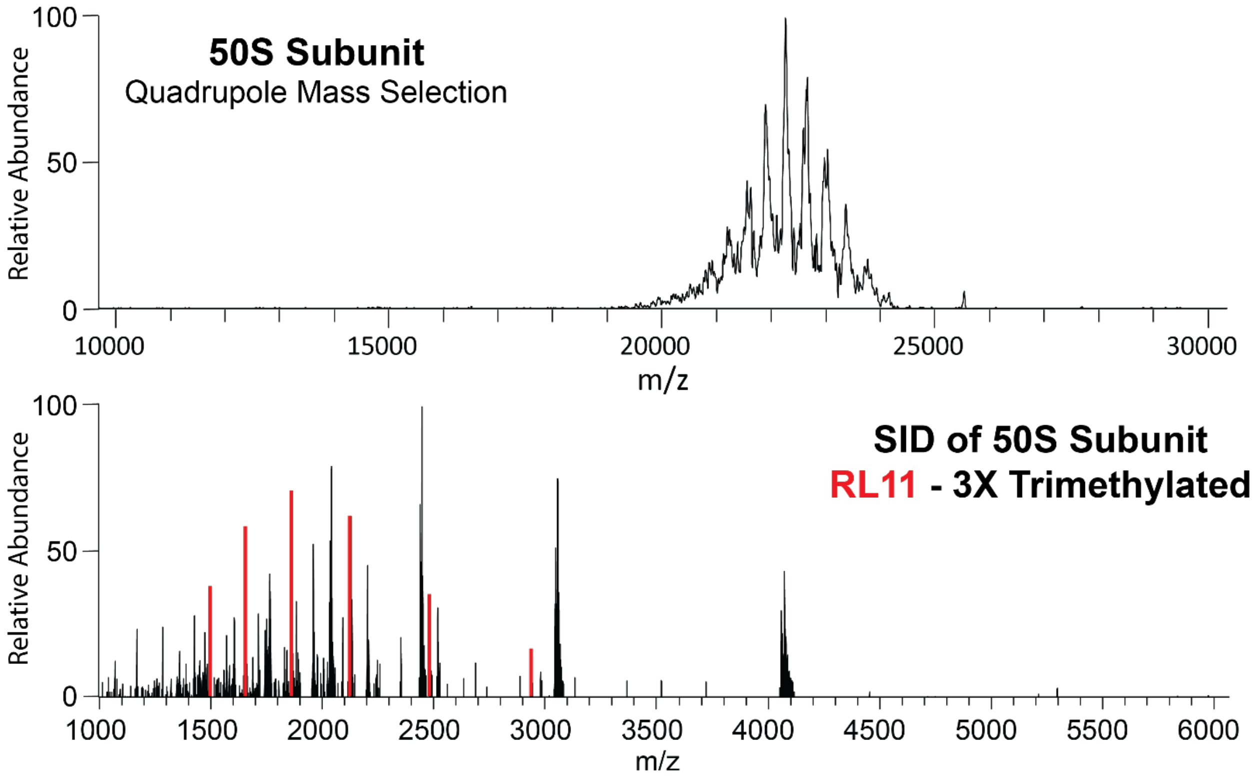


**Figure S1.** Quadrupole mass selection of E. coli ribosome 50S subunit (top) and SID of the 50S subunit (bottom). The SID feature of the combination ExD-SID device yielded abundant ribosomal protein subunits with, for example, 3x trimethylated RL11 measured at 14,861.0 ± 0.5 ppm (charge state distribution marked in red).

**Table S1**. Typical voltages applied to the ExD cell for electron capture charge reduction.  The ExD-SID cell of Figure 1 is repeated here for comparison with the descriptors in the Table.

| | ECCR 2V | ECCR 4V | Maximal ECCR |
|---|---|---|---|
| **Entrance quadrupole** | 0 V | 0 V | 0 V |
| **Entrance lens** | -30 V | -30 V | -30 V |
| **Magnet lens 1** | 2 V | 4 V | 17 V |
| **Filament holder** | 3 V | 5 V | 17 V |
| **Filament Bias** | -0.5 V | -0.5 V | -1.5 V |
| **Magnet lens 2** | 2 V | 4 V | 17 V |
| **Exit quadrupole** | -6 V | -6 V | -6 V |
| **Exit lens** | -30 V | -30 V | -30 V |
| **Filament current** | 2.2 A | 2.2 A | 2.2 A |
| **Surface** | -8 V | -8 V | -8 V |
| **Extractor** | -8 V | -8 V | -8 V |
| **Deflector** | -8 V | -8 V | -8 V |

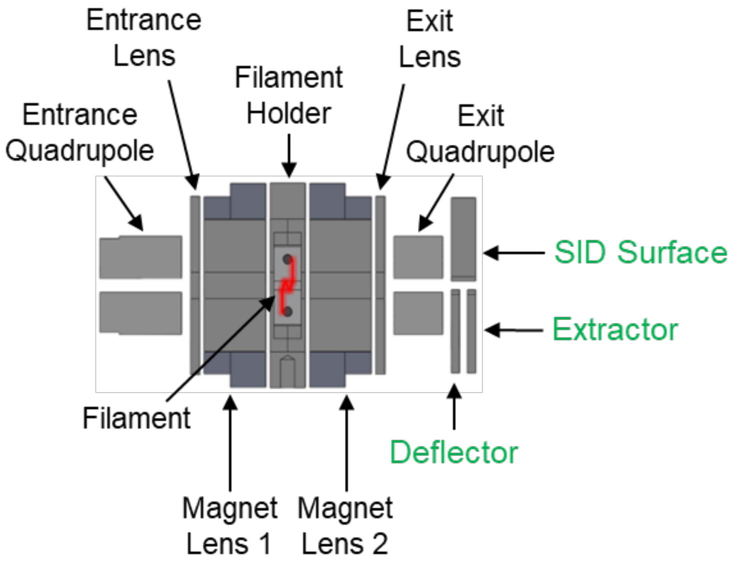

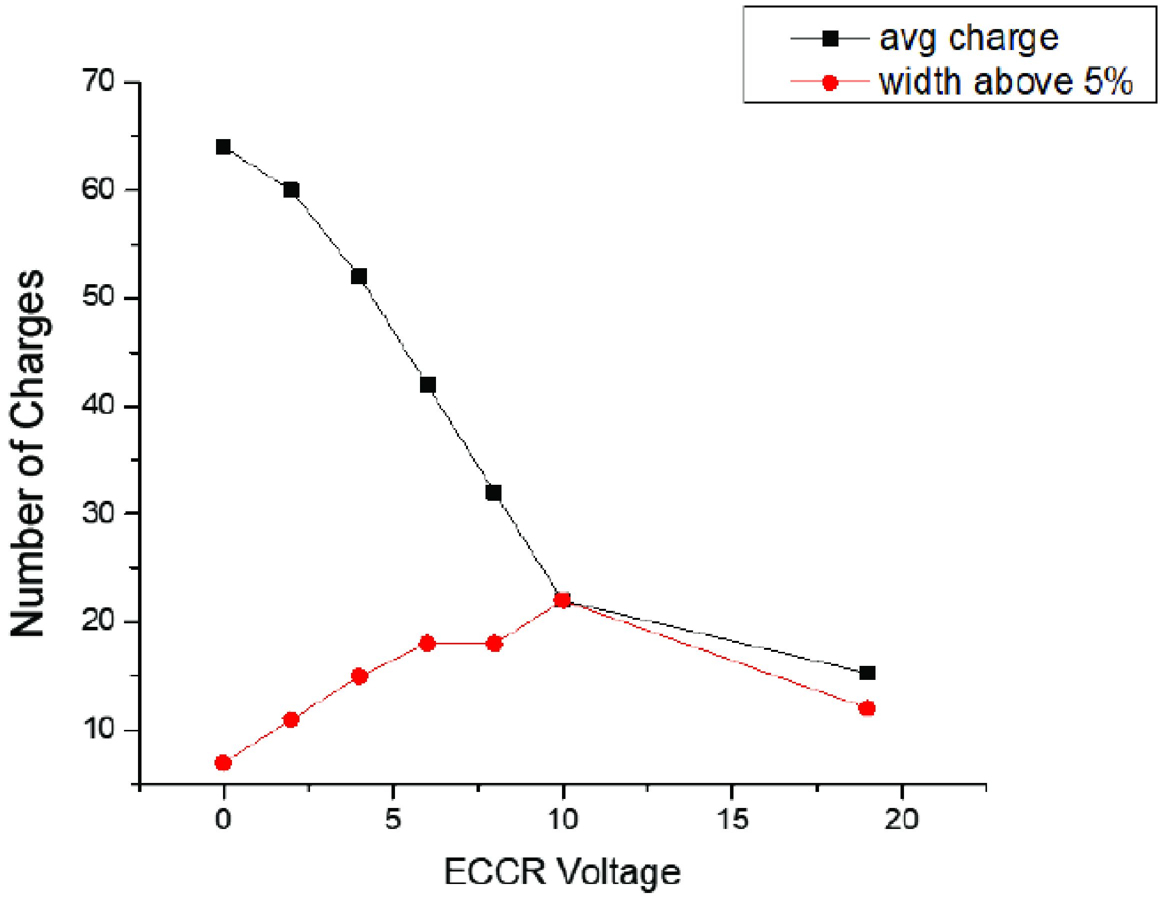


**Figure S2**. Average charge of GroEL and width of charge state distribution for peaks above 5% relative intensity as a function of ECCR voltage. ECCR voltage, here, refers to the potential applied to magnet lens 1 and 2 which is increased (along with the filament holder voltage) to increase the extent of ECCR. Voltages applied to magnetic lens 1 and 2 and the filament holder were changed in 2 V steps over the ECCR voltage range of 2-10 V, representative settings for ECCR 2V and 4V are shown in Table S1. The last point was obtained at maximal ECCR, the settings for which are shown in Table S1.

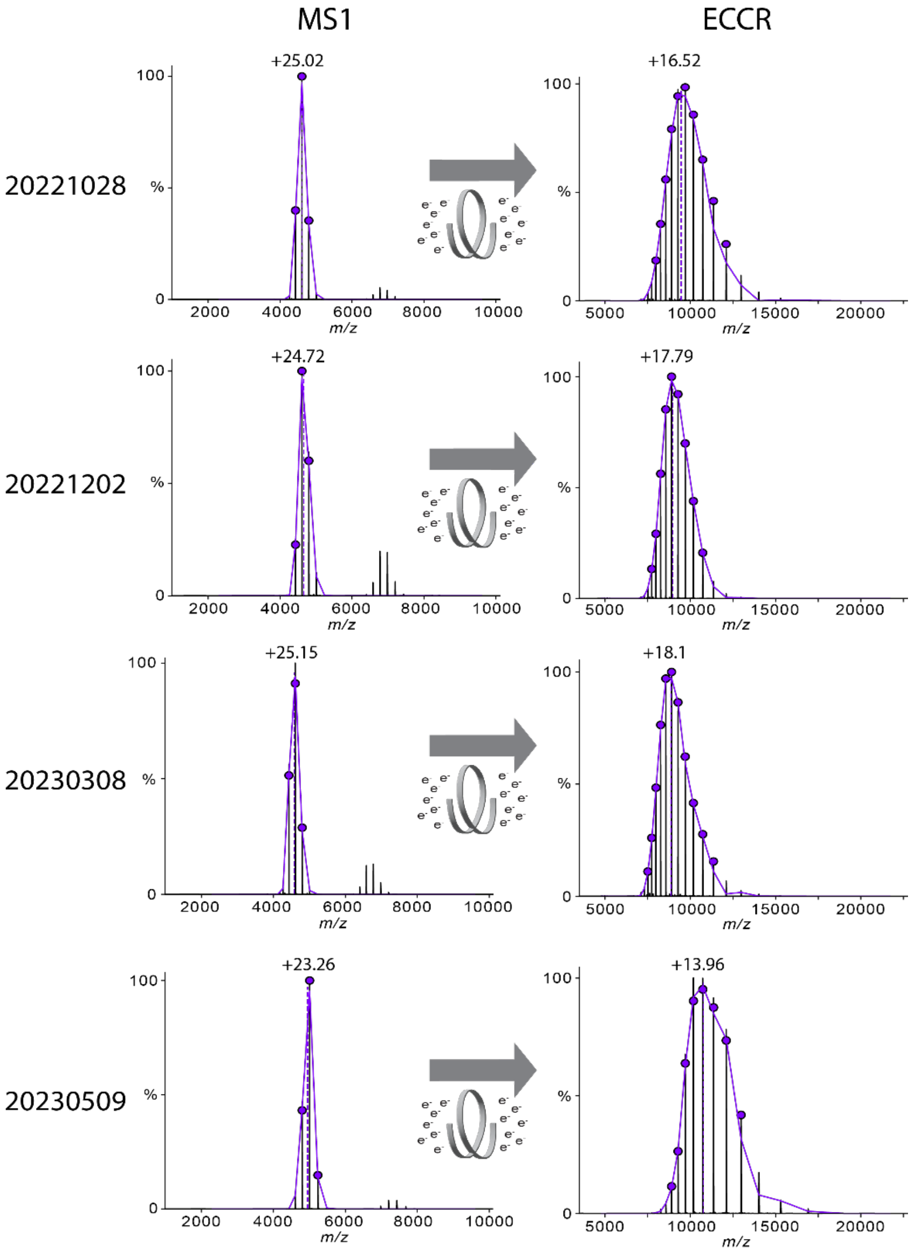


**Figure S3**. Long term stability and reproducibility of ECCR for C-Reactive Protein pentamer. The charge state distribution from *m/z* 4000 to 6000 was quadrupole mass selected and subjected to ECCR using the same voltage profile over 7 months. MS1 and ECCR charge state distributions are labeled with the weighted average charge. Replicates are shown over 7 months of use, and the average charge state before ECCR is +24.5 ± 0.9, while the average charge state after ECCR is +16.6 ± 1.9 (± refers to standard deviation). The voltages applied to the ExD-SID device are shown in Table S2. (Calendar dates of acquisition are shown on the left in year-month-day format.)

**Table S2**. Voltage profiles applied to the ExD-SID device for C-Reactive Protein Flythrough (MS1) and ECCR.

| | Entrance Quadrupole (V) | Entrance Lens (V) | Magnet Lens 1 (V) | Filament Holder (V) | Filament Bias (V) | Magnet Lens 2 (V) | Exit Quadrupole (V) | Exit Lens (V) |
|---|---|---|---|---|---|---|---|---|
| Flythrough | 0 | -19.0 | 0.0 | -2.0 | -3.5 | 0.0 | -0.5 | -19.0 |
| ECCR | 0 | -30.0 | 7.0 | 8.0 | -0.5 | 7.0 | -5.0 | -30.0 |

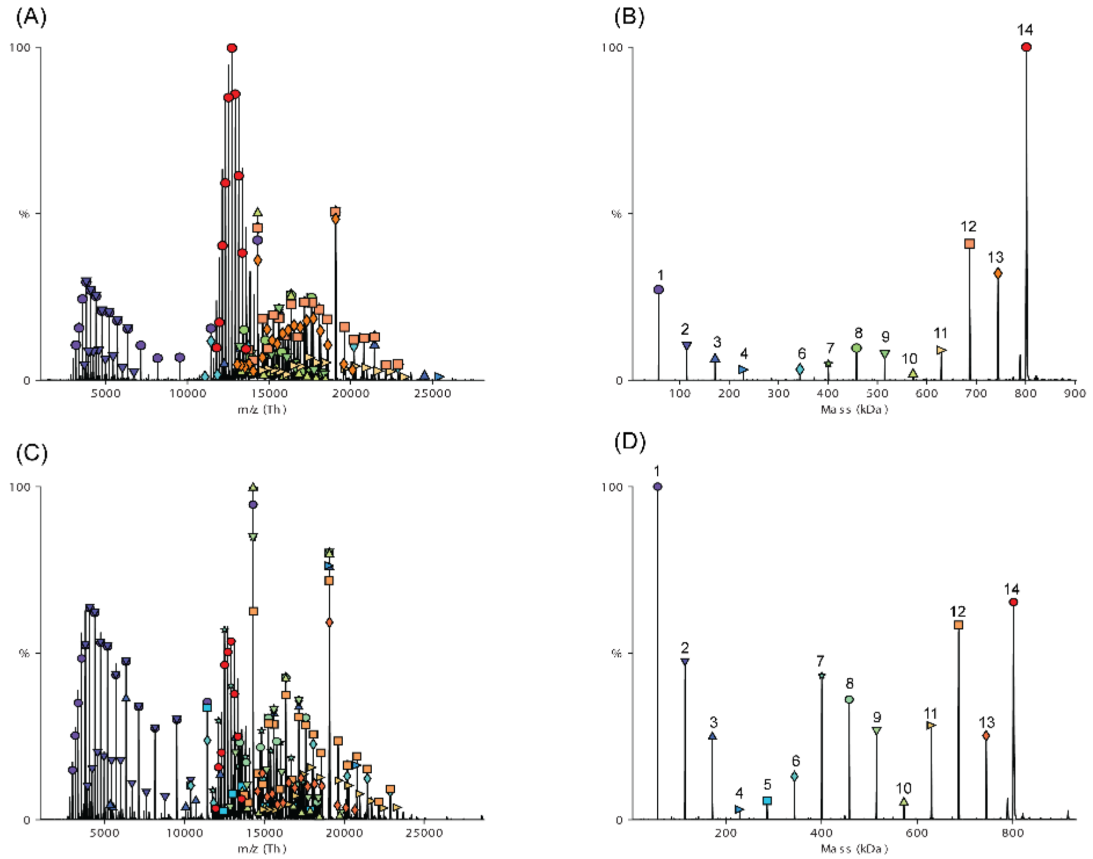


**Figure S4.** GroEL SID with normal charge (68+ weighted average charge). (A) raw data at SID 125 V; (B) deconvolved data at SID 125 V using UniDec.[48] (C) raw data at SID 145 V; (D) deconvolved data at SID 145 V using UniDec.[48]

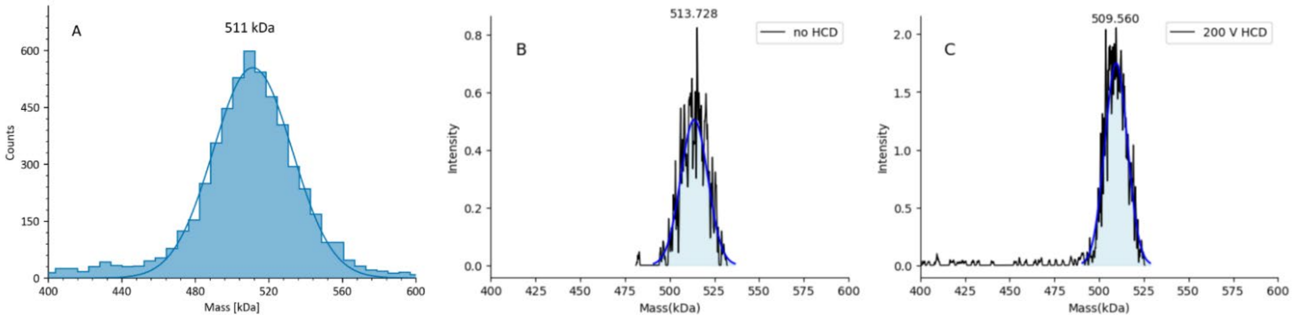


**Figure S5.** A) Mass photometry (Refeyn TwoMP) of VFLIP spike protein trimer, showing the average mass of 510 kDa ± 5%. Charge detection mass spectrometry of heterogeneous VFLIP spike protein with an in-source trapping voltage of -50 V and B) no HCD or C) 200 V HCD. HCD removes some salt and/or solvent adducts, but we cannot rule out minor covalent losses.

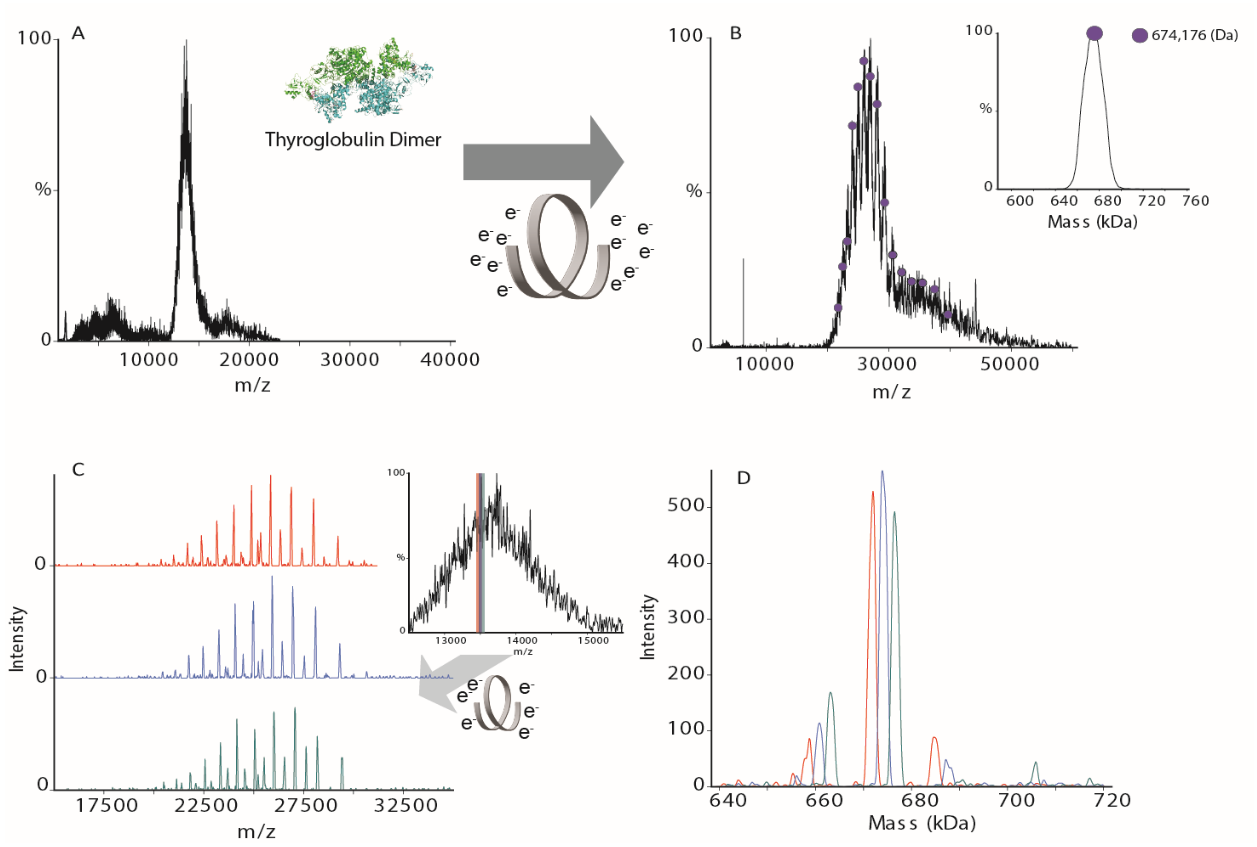


**Figure S6.** A) An unresolved native mass spectrum of dimeric bovine thyroblobulin, Tg. PDB: 7QTQ (B) A charge state-resolved native mass spectrum of the bovine TG with electron capture charge reduction (ECCR, voltage 7 V). The isolation region is 12k-17.5k *m/z*. The average mass is 674 kDa deconvolved using UniDec.[48] (C) Overlay of three narrow window isolations (each highlighted in a different color) covering the *m/z* range of 13426-13599, windows share an overlap of 1 *m/z*, inset showing the position of three narrow quadrupole window selections. (D) Deconvolved mass spectrum of the three narrow window isolations shown in C, deconvolved using UniDec.[48]

To further assess the ability of ECCR to provide insight into heterogeneous glycoproteins we also considered the glycoprotein thyroglobulin. Thyroglobulin (Tg) is a dimeric protein complex that exhibits multiple PTMs, including glycosylation, phosphorylation, and iodinated tyrosine. While both chains in the dimer have the same sequence of amino acids, they may exhibit differences in their PTMs. The sequence mass of bovine thyroglobulin is ~602 kDa, however, it is commonly referred to as a dimer of ~670 kDa partially because of extensive glycosylation (~10% of its mass being attributed to glycosylation). When Tg is introduced into the mass spectrometer under native like conditions, like with VFLIP, it is not possible to resolve the charge states and determine the mass (Figure S6A). However, after gas-phase charge reduction of the entire charge state envelope, an average mass of 674 kDa was determined. As with VFLIP, when narrow window isolation is used, the complexity and heterogeneity of the sample becomes more apparent, as shown in Figures S6C and D. **Because Tg is often used as a mass photometry calibrant, we recommend ECCR mass measurement of a user's chosen Tg calibrant to determine the mass used in the MP calibration curve.**

**METHODS**

***Materials and Sample Preparation.***

C-reactive protein, GroEL, β-Amylase (BAM), and thyroglobulin were purchased from Sigma Aldrich. VFLIP (a stable, covalently linked SARS-coV-2 spike trimer with the D614G mutation of the Wuhan variant) was expressed and purified as previously described.[1] GroEL was refolded based on a previously described protocol.[2] CRP, thyroglobulin, GroEL and VFLIP were buffer exchanged into 200 mM ammonium acetate using BioRad micro P6 columns. VFLIP for narrow window selection was buffer exchanged into 200 mM ammonium acetate using 50 kDa molecular weight cut-off (MWCO) Amicon® Ultra Centrifugal Filters (Sigma Aldrich). Samples were run at approximately 1 µM complex concentration for native mass spectrometry, 500 nM for CDMS, and 10 nM for mass photometry.

***Mass Spectrometry.***

All experiments were performed using a Q Exactive UHMR Orbitrap mass spectrometer (Thermo Fisher Scientific, Bremen, Germany) fitted with a hybrid device enabling electron-based fragmentation and surface induced dissociation, here referred to as the ExD-SID cell (cell generated by modifying the ExD cell from e-MSion Inc., Corvallis, OR with SID, see results). The ExD-SID cell replaced the transfer multipole between the quadrupole mass filter and C-trap as previously described for the standard ExD cell[3–5] and standard 4 cm SID cell.[6]

Static nanoelectrospray ionization was performed using uncoated glass capillary emitters pulled in-house from borosilicate glass (O.D 1mm I.D 0.78 mm) with a filament using a Sutter P-97 micropipette puller. A platinum wire was inserted into the emitter, and the electrospray voltage was directly applied to the analyte solution. Typical electrospray voltage was 900 V. In-source trapping collision energy for desolvation was optimized for each sample and was generally between -30 and -100 V. An inlet capillary temperature of 250 °C, S-lens RF of 200, and a fixed ion injection time of 50−1000 ms were used for all experiments. Approximately 2-5 minutes of averaging was used to produce ECCR, SID, and ECCR-SID spectra. For narrow window selection experiments approximately 6 minutes of averaging was performed.

Deconvolution and average charge state analysis was performed using UniDec.[7] The UniDec parameters for Figure 5C are as follows; Charge Range: 1-60, Mass range 400-600 kDa, Sample Mass Every (Da) 1, Peak FWHM(Th) 200.0, Beta: 50.0, Charge Smooth Width: 1.0, Point Smooth Width: 10.0, Mass Smooth Width: 0.0, Maximum # of Iterations: 100, Peak Detection 5000 Da, Peak Detection Threshold 0.5. For the rest narrow quadrupole selections, the parameters were Charge Range: 1-50, Mass range 400-600 kDa, Sample Mass Every (Da) 1, Peak FWHM(Th) 5.0, Beta: 0, Charge Smooth Width: 1.0, Point Smooth Width: 1, Mass Smooth Width: 0, Maximum # of Iterations: 100, Peak Detection 500 Da, Peak Detection Threshold 0.1. The CDMS data were acquired at 200,000 resolving power at *m/z* 400 and trapping gas 2 using Direct Mass Technology mode. The CDMS datasets were processed and deconvolved by STORIboard software (Thermo Fisher Scientific). We note here that with extensive use of the ExD-SID cell, some surface contamination is detected; work is in progress to define the problem and resolve it with a cell redesign, although we continue using the device.

***Monte Carlo simulation***

To determine the theoretical masses of the glycosylated trimer, a Monte Carlo simulation was performed by using the glycan composition and corresponding occupancy percentage from glycoproteomics papers[8,9], which provided data for 20 N-glycan sites and 8 O-glycan sites. N-glycan site occupancies were determined by comparing the peak intensities of identified glycopeptide and unoccupied peptide with normalization.[8] O-glycan site occupancies were quantified by comparison of precursor ion intensities of unoccupied peptides and deglycosylated peptides following Peptide: N-glycosidase F (PNGase F) de-glycosylation under $^{18}O$ water, as Sanda and coworkers described previously.[9,10] For theoretical mass determination of the spike trimer, a random glycan composition was chosen, from the known glycosylation states, using a homemade Python program for each glycan site to calculate the total glycan mass for the monomer of the spike protein and the possibility of choosing any glycan composition was specified using the occupancy percentage for each glycan composition. Then, three monomer masses and three randomly selected total glycan masses were summed to calculate the total trimer mass. The equation for calculating one possible total trimer mass is listed below:

$$trimer\ mass = monomer\ mass\ \times 3 + (total\ glycan\ mass\ 1) + (total\ glycan\ mass\ 2) + (total\ glycan\ mass\ 2)$$

Total glycan mass 1, total glycan mass 2, and total glycan mass 3 are the sum of the random selection for all 28 glycan sites, resulting in possible different glycan masses in monomer for a trimer spike protein. By repeating this process 7.5 $\times 10^6$ times, a list of the possible total trimer masses was obtained. The final trimer masses were rounded into integers for faster data analysis.

***Mass photometry (MP)***

A Refeyn TwoMP was used for MP experiments. HybriSlip™ Hybridization Covers and CultureWell™ Reusable gaskets (Grace Biolab) were cleaned using water and isopropanol. MP was calibrated using a mixture of β-Amylase (monomer 56 kDa, dimer 112 kDa, tetramer 224 kDa) and GroEL (tetradecamer 801 kDa) in 200 mM ammonium acetate solution. All data were collected using AcquireMP software for one minute. The data were processed using DiscoverMP software.

**References**


(1) Olmedillas, E.; Mann, C. J.; Peng, W.; Wang, Y.-T.; Avalos, R. D.; Bedinger, D.; Valentine, K.; Shafee, N.; Schendel, S. L.; Yuan, M.; Lang, G.; Rouet, R.; Christ, D.; Jiang, W.; Wilson, I. A.; Germann, T.; Shresta, S.; Snijder, J.; Saphire, E. O. Structure-Based Design of a Highly Stable, Covalently-Linked SARS-CoV-2 Spike Trimer with Improved Structural Properties and Immunogenicity. bioRxiv May 6, 2021, p 2021.05.06.441046. https://doi.org/10.1101/2021.05.06.441046.

(2) Zhou, M.; Jones, C. M.; Wysocki, V. H. Dissecting the Large Noncovalent Protein Complex GroEL with Surface-Induced Dissociation and Ion Mobility–Mass Spectrometry. *Anal. Chem.* **2013**, *85* (17), 8262–8267. https://doi.org/10.1021/ac401497c.

(3) Zhou, M.; Liu, W.; Shaw, J. B. Charge Movement and Structural Changes in the Gas-Phase Unfolding of Multimeric Protein Complexes Captured by Native Top-Down Mass Spectrometry. *Anal. Chem.* **2020**, *92* (2), 1788–1795. https://doi.org/10.1021/acs.analchem.9b03469.

(4) Shaw, J. B.; Malhan, N.; Vasil'ev, Y. V.; Lopez, N. I.; Makarov, A.; Beckman, J. S.; Voinov, V. G. Sequencing Grade Tandem Mass Spectrometry for Top–Down Proteomics Using Hybrid Electron Capture Dissociation Methods in a Benchtop Orbitrap Mass Spectrometer. *Anal. Chem.* **2018**, *90* (18), 10819–10827. https://doi.org/10.1021/acs.analchem.8b01901.

(5) Shaw, J. B.; Liu, W.; Vasil'ev, Y. V.; Bracken, C. C.; Malhan, N.; Guthals, A.; Beckman, J. S.; Voinov, V. G. Direct Determination of Antibody Chain Pairing by Top-down and Middle-down Mass Spectrometry Using Electron Capture Dissociation and Ultraviolet Photodissociation. *Anal. Chem.* **2020**, *92* (1), 766–773. https://doi.org/10.1021/acs.analchem.9b03129.

(6) VanAernum, Z. L.; Gilbert, J. D.; Belov, M. E.; Makarov, A. A.; Horning, S. R.; Wysocki, V. H. Surface-Induced Dissociation of Noncovalent Protein Complexes in an Extended Mass Range Orbitrap Mass Spectrometer. *Anal. Chem.* **2019**, *91* (5), 3611–3618. https://doi.org/10.1021/acs.analchem.8b05605.

(7) Marty, M. T.; Baldwin, A. J.; Marklund, E. G.; Hochberg, G. K. A.; Benesch, J. L. P.; Robinson, C. V. Bayesian Deconvolution of Mass and Ion Mobility Spectra: From Binary Interactions to Polydisperse Ensembles. *Anal. Chem.* **2015**, *87* (8), 4370–4376. https://doi.org/10.1021/acs.analchem.5b00140.

(8) Wang, D.; Zhou, B.; Keppel, T. R.; Solano, M.; Baudys, J.; Goldstein, J.; Finn, M. G.; Fan, X.; Chapman, A. P.; Bundy, J. L.; Woolfitt, A. R.; Osman, S. H.; Pirkle, J. L.; Wentworth, D. E.; Barr, J. R. N-Glycosylation Profiles of the SARS-CoV-2 Spike D614G Mutant and Its Ancestral Protein Characterized by Advanced Mass Spectrometry. *Sci. Rep.* **2021**, *11* (1), 23561. https://doi.org/10.1038/s41598-021-02904-w.

(9) Sanda, M.; Morrison, L.; Goldman, R. N- and O-Glycosylation of the SARS-CoV-2 Spike Protein. *Anal. Chem.* **2021**, *93* (4), 2003–2009. https://doi.org/10.1021/acs.analchem.0c03173.

(10) Pompach, P.; Brnakova, Z.; Sanda, M.; Wu, J.; Edwards, N.; Goldman, R. Site-Specific Glycoforms of Haptoglobin in Liver Cirrhosis and Hepatocellular Carcinoma *. *Mol. Cell. Proteomics* **2013**, *12* (5), 1281–1293. https://doi.org/10.1074/mcp.M112.023259.
